# Detection and Removal of Hyper-synchronous Artifacts in Massively Parallel Spike Recordings

**DOI:** 10.1101/2024.01.11.575181

**Authors:** Jonas Oberste-Frielinghaus, Aitor Morales-Gregorio, Simon Essink, Alexander Kleinjohann, Cristiano A. Köhler, Frederic Barthélemy, Alexa Riehle, Thomas Brochier, Simon Musall, Junji Ito, Sonja Grün

**Affiliations:** Institute for Advanced Simulation (IAS-6), Jülich Research Centre, Jülich, Germany; RWTH Aachen University, Aachen, Germany; Institut de Neurosciences de la Timone (INT), Aix-Marseille Université, CNRS, Marseille, France; Institute of Biological Information Processing (IBI-3), Jülich Research Centre, Jülich, Germany; Institute of Biology 2, RWTH Aachen University, Aachen, Germany; Faculty of Medicine, Institute of Experimental Epileptology and Cognition Research, University of Bonn, Bonn, Germany; JARA Brain Institute I (INM-10), Jülich Research Centre, Jülich, Germany; Theoretical Systems Neurobiology, RWTH Aachen University, Aachen, Germany

**Keywords:** artifacts, electrophysiology, Neuropixel, preprocessing, spikes, Utah arrays

## Abstract

Contemporary electrophysiology experiments often involve massively parallel recordings of neuronal activity using multi-electrode arrays. While researchers have been aware of artifacts arising from electric cross-talk between channels in setups for such recordings, systematic and quantitative assessment of the effects of those artifacts on the data quality has never been reported. Here we present, based on examination of electrophysiology recordings from multiple laboratories, that multi-electrode recordings of spiking activity commonly contain extremely precise (at the data sampling resolution) spike coincidences far above the chance level. We derive, through modeling of the electric cross-talk, a systematic relation between the amount of such hyper-synchronous events (HSEs) in channel pairs and the correlation between the raw signals of those channels in the multi-unit activity frequency range (500-7500 Hz). We show that whitening the band-pass filtered raw signals removes the above chance HSEs; strongly suggesting they originate from linear mixing of signals. Whitening should therefore be performed prior to spike sorting and any further analysis of precise spike correlation, otherwise analysis results may be considerably affected.

**Significance Statement:** Artifacts are ubiquitous in electrophysiological recordings. To mitigate their impact, these artifacts need to be detected and they should be removed from the data without impacting the quality of the data. This work presents measures to identify and quantify the amount of artifacts within a multichannel recording by evaluating the occurrence of hyper-synchronous events i.e., spikes that are synchronous on a sub-millisecond time scale, and further introduces zero-phase component analysis (ZCA) as a method to remove these artifacts from the data. Thus, we recommend to use ZCA as a general preprocessing for electrophysiological recordings.

## Introduction

Modern electrophysiological experiments often use multielectrodes to record simultaneously many neurons from multiple sites. These recordings are known to be very sensitive to various sources of noise (Rey et al., 2015; Harris et al., 2016), which need to be mitigated to enable reliable and robust data analysis (e.g., (Oberste-Frielinghaus et al., 2025)). Artifacts in neural recordings may occur from both, internal (heartbeats, eye movements, chewing, muscles, etc., (Merletti and Conte, 1997)), and external (electric grid interference, loud noises etc.) sources (Fabietti et al., 2020). These sources are of non-neural origin, and can often be distinguished from neural activity by their particular frequencies and distinct waveforms in the raw signals. This kind of noise can be removed by applying high-pass filtering to the signals and/or by discarding the data during the time periods, e.g. of body movements. In more general cases, more sophisticated methods are required, such as independent component analysis (Bell and Sejnowski, 1995; Rong and Contreras-Vidal, 2006; Delorme et al., 2007), canonical correlation analysis (Hotelling, 1936; Wim De Clercq et al., 2006), wavelet methods, or certain combinations thereof. These are mostly applied to electroencephalogram (EEG) (Wim De Clercq et al., 2006; Delorme et al., 2007; Shackman et al., 2009; Barban et al., 2021; Mumtaz et al., 2021) and magnetoencephalogram (Rong and Contreras-Vidal, 2006) data, and to a lesser extent to extracellular electrophysiology (Ludwig et al., 2009; Islam et al., 2014; Fabietti et al., 2020). Besides these common noise sources, (capacitive, resistive, or inductive) coupling between shanks or cables can cause cross-talk between different channels (Nelson et al., 2017), leading to a mixing of neural signals that can persist even after spike sorting (Torre et al., 2016). Cross-talk is a well-known problem in EEG (Nagaoka et al., 1992) and electromyogram (Koh and Grabiner, 1993; Kilner et al., 2002; Farina et al., 2004) data recording systems, however only a few studies have focused on cross-talk in electrophysiology systems (Musial et al., 2002; Nelson et al., 2017; Pérez-Prieto and Delgado-Restituto, 2021). One approach to avoid cross-talk in these systems is through hardware implementations, like improving the isolation and design of the circuits (Blot and Barbour, 2014; Nelson et al., 2017; Pérez-Prieto and Delgado-Restituto, 2021; Perez-Prieto et al., 2021). However, even with the best efforts, eliminating all possible causes of artifacts is practically unfeasible. Thus, the recorded data need to be examined offline to check for existence of artifacts, and if they exist, they need to be removed from the data, otherwise they lead to false results, especially in the context of analysis of fine temporal correlations in spike trains (Oberste-Frielinghaus et al., 2025).

We here illustrate, on three different data sets from three different laboratories, animals and recording setups, hyper-synchronous artifacts that are present with a time resolution of the recording system (30kHz). Two of the data sets were recorded by Utah arrays (Blackrock Microsystems, Salt Lake City, UT, USA, (Hatsopoulos et al., 1998; de Haan et al., 2018; Chen et al., 2022) and the third by a Neuropixel probe (IMEC, Leuven, Belgium sold by Cambridge NeuroTec) (Jun et al., 2017). Both systems have very different approaches to measure single neuron contributions. The electrodes on Utah arrays are at least 400 µm apart such that the same single neuron cannot be identified on different electrodes, while in Neuropixel probes the recordings of a single neuron on neighboring electrodes (inter-contact distance of 20 µm is intended. The extraction of single units is in both cases performed by spike sorting after some preprocessing (high pass filtering, etc.) of the raw recorded signals. Still, in data of both recording types we identified hypersynchronous artifacts even after application of spike sorting. Thus the artifacts appear as extremely precise spike synchrony (at the data sampling resolution, e.g 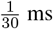) (Yu et al., 2009; Torre et al., 2016; Dehnen et al., 2021; Chen et al., 2022), which we identified as non-neuronal.

After demonstrating on these three data sets how hyper-synchronous artifacts or their derivations are identified, we introduce a software solution to get rid of them. We illustrate that the artifacts can be removed by using the zero-phase component analysis (ZCA) whitening (Bell and Sejnowski, 1997; Kessy et al., 2018). This method is often part of spike sorting software solutions (e.g., Mountainsort, Kilosort), but there it is only applied locally on a subset of the channels. We show that it should be applied to all recordings channels together to secure removal of artifacts.

## Materials and Methods

### Experimental Design

*Macaques* We analyzed resting state data from two (N=2) rhesus macaques (*Macaca mulatta*), recorded in two different experimental laboratories. The data from macaque L was collected at the Netherlands Institute for Neuroscience, and previously published (Chen et al., 2022). The data from macaque Y was collected at the Institut de Neurosciences de la Timone, with the recording apparatus described by de Haan et al. (2018). At the time of the array implantation macaque L (male) was 7 years old and weighed ∼ 11 kg; and macaque Y (female) was 6 years old and weighed ∼ 7 kg. All experimental and surgical procedures for macaque L complied with the NIH Guide for Care and Use of Laboratory Animals, and were approved by the institutional animal care and use committee of the Royal Netherlands Academy of Arts and Sciences (approval number AVD-8010020171046). All experimental and surgical procedures for macaque Y were approved by the local ethical committee (C2EA 71; authorization Apafis#13894-2018030217116218v4) and conformed to the European and French government regulations.

*Mouse* Mouse data were recorded at the Institute of Biology 2 at RWTH Aachen using the recording apparatus described by Balla et al. (2023). The mouse was an 18-week-old C57BL/6J animal and weighed 26 g at the time of the acute recording. All procedures for the mouse experiment were performed in compliance with the EU directives 86/609/EWG and 2007/526/EG guidelines for animal experiments and were approved by the local governments (Thueringer Landesamt, Bad Langensalza, Germany and Landesamt für Natur, Umwelt und Verbraucherschutz Nordrhein-Westfalen, Recklinghausen, Germany) under permit number 81-08.04.2021.A021.

### Data collection

Both macaques were recorded using setups based on the Blackrock Microsystems (Blackrock Neurotech, Salt Lake City, USA) ecosystem.

*Macaque L* See Chen et al. (2022) for an in-depth description of this data set. Briefly, electrophysiological signals originating from the V1 and V4 regions were recorded using a configuration of 1024 channels distributed across 16 Utah Arrays, each comprising 8x8 electrodes. The references wires were located on alternating arrays (array numbers 1, 3, 5, 7, 9, 11, 13, and 15) and ran alongside the wire bundle before emerging several millimetres before the point of connection between the wire bundle and the array. Sampling was performed at a rate of 30 kHz. The signal pathway commenced with the passive conduction of neuronal signals from the 1024-channel pedestal to an Electronic Interface Board (EIB). This EIB was equipped with 32 36-channel Omnetics connectors, facilitating the interface with eight 128-channel CerePlex M headstages. Each CerePlex M headstage received signals from two electrode arrays and applied a 0.3–7500 Hz analog filter at unity gain, refraining from signal amplification. Analog-to-digital conversion (ADC) was conducted by the CerePlex M, employing a 16-bit resolution with a sensitivity of 250 nV/bit. The digitized signal was subsequently routed to a 128-channel Digital Hub, with each Digital Hub processing data originating from one CerePlex M. The Digital Hub undertook the conversion of the digital signal into an optic-digital format. This transformed signal was then transmitted via an optic-fiber cable to a 128-channel Neural Signal Processor (NSP) for subsequent processing and storage. Each Digital Hub supplied the signal to a single NSP. In total, eight NSPs were employed. In the present study, we focused on a single data stream (NSP1) from one session, which included one array in V1 and one array in V4. These arrays showed many artifacts in a previous less exhaustive analysis (Chen et al., 2022), including events across both arrays, even though they were separated by the lunate sulcus. Thus, this data stream was particularly useful to demonstrate the non-neural origin of the artifacts. We manually found a large artifact event (see Common noise artifact example) which we cut out before the analysis of the data.

*Macaque Y* Electrophysiological signals originating from V1, V2, DP, and 7A were recorded with four separate 6x6-electrode Utah Arrays. Additionally, M1/PMd was recorded with a 10x10-electrode Utah Array. Sampling was performed at a rate of 30 kHz. In contrast to macaque L, the signal pathway commenced with two 128-channel pedestals (CerePort): One for the four 36-channel arrays and one for the 100-channel array. There are two reference electrodes per pedestal consisting in silver wires placed on the dura at two different sites under the skull. Each of the pedestals interfaced with a 128-channel CerePlex E headstage. One of the two reference electrodes can be selected with a microswitch on the Cereplex. When the animal recovered from surgery and during the first recording session, both references were tested and the one that gave the best signal to noise ratio was selected. Once it was chosen, the reference was kept for the sake of consistency across the recording session. Notably, the CerePlex E headstages applied a 0.3–7500 Hz analog filter at unity gain, refraining from signal amplification. Analog-to-digital conversion (ADC) was conducted by the CerePlex E, employing a 16-bit resolution with a sensitivity of 250 nV/bit. The digitized signal was subsequently routed to a 128-channel Digital Hub, with each Digital Hub processing data originating from one CerePlex E, which in turn was linked to one 100-channel array or four 36-channel arrays. The Digital Hub undertook the conversion of the digital signal into an optic-digital format. This transformed signal was then transmitted via an optic-fiber cable to a 128-channel Neural Signal Processor (NSP) for subsequent processing and storage. Each Digital Hub supplied the signal to a single NSP. In total, two NSPs were employed. In the present study, we focused on the M1/PMd array from one session, to demonstrate that the artifacts are not limited to visual processing areas.

*Mouse* High-density electrophysiological recordings were done with Neuropixels 1.0 probes in awake head-fixed mice, freely running on a wheel. To allow a later recovery of the probe position in the brain, the probe was painted with DiD cell labeling solution (Invitrogen V22887). High pass-filtered data above 300 Hz at 30 kHz and low pass-filtered signals between 0.5-100 Hz at 2.5 kHz from the bottom 384 channels of the Neuropixels probe (∼3.8 mm active recording area) were recorded. To simultaneously record from V1 and SC, the probes were angled to 35° elevation and inserted to a depth of 3.6 mm. The reference electrode was a gold-pin and it was placed on the dura above the cerebellum. Visual stimuli were presented during the recording, consisting of 0.1-s long full-field low-pass filtered Gaussian noise patterns with a cutoff at 0.12 cycles per degree and a temporal frequency of 1 Hz. Each stimulus was presented twice with an inter-trial interval of 3-5 seconds. Signals were acquired with an external Neuropixels PXIe card (IMEC, Belgium) used in a PXIe-Chassis (PXIe-1071, National Instruments, USA). Triggers and control signals for different stimuli and wheel movements were separately recorded as analog and digital signals using the SpikeGLX software (Janelia Farm Research Campus, USA; Bill Karsh, https://github.com/billkarsh/SpikeGLX).

### Signal processing

*Macaque recordings* All offline signal processing steps (except for spike sorting) and data analysis described below were performed by original custom codes written in the Python programming language and executed in a pipeline defined as a Snakemake workflow (Mölder et al., 2021). The recorded raw signals were band-pass filtered in the frequency range of 250 Hz 7500 Hz − (second order Butterworth filter, as implemented in the `signal_processing.butter()` function in the Elephant toolbox (Denker et al., 2018)). According to Quiroga (2007), putative spikes were extracted from the band-pass filtered raw signal of each channel by thresholding it at

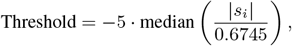

where *s*_*i*_ denotes the band-pass filtered raw signal of channel *i*. The time of threshold crossing of each putative spike was registered as the time of that spike. The resulting spike times per channel were taken as the spike train of the multi-unit activity of that channel. For the artifact removal, we apply ZCA whitening after the band-pass filter and then proceed as above.

*Mouse recording* The data was processed with the SpikeInterface toolbox (Buccino et al., 2020). First, the data were band-pass filtered between 500 and 7500 Hz. Second, the channels that were broken or outside the brain were removed from further analysis, using a manually curated bad channels rejection implementation. Third, the data were time-shifted to correct for multiplexing during acquisition. Fourth, we performed a median-subtraction across all channels, to remove the common signal originating from the reference electrode. Finally, the data were automatically spike-sorted using Kilosort 4.0.6 (Pachitariu et al., 2024). Three different parameter settings were used for the ZCA whitening step before spike sorting: 1) without whitening; 2) whitening only the nearest 32 channels to each electrode (default setting of Kilosort); and 3) whitening across all channels. All other parameters were left unchanged from their default values.

### Zero-phase component analysis whitening

As a data cleaning method for removing correlations introduced by the electric cross-talks between channels, we propose whitening of the signals by use of zero-phase component analysis (ZCA) (Bell and Sejnowski, 1997; Kessy et al., 2018). ZCA is the whitening transformation of given multivariate data that is closest to the original data in terms of Euclidean distance. For an *N*_channels_ ×*N*_samples_ data matrix *X*, the ZCA transform matrix *W* is defined using the covariance matrix Σ of *X* as *W* = Σ^−1*/*2^. Then the whitened data 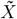 are obtained by applying this transform matrix *W* to the original data matrix *X* as 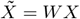. For numerical stability, in practical applications to the experimental datasets the transform matrix *W* was computed as *W* = *U* (Λ+ *ϵI*)^−1*/*2^*U* ^*T*^, with *U* and Λ obtained from the eigendecomposition of the covariance matrix: Σ = *U* Λ*U* ^*T*^ and *ϵ* set to 10^−5^.

### Cross-talk model

We observed spike waveforms that are very similar among multiple channels without time delay or phase shift (Figure 2a). This led us to assume the cross-talk between channels to be a resistive interaction, which is instantaneous and has no frequency dependence, rather than a capacitive or inductive interaction, which causes frequency dependent amplitude reduction and phase shift. For modeling the generation of artifact spikes in a pair of channels, we formulated this assumption as a linear mixture of signals between channels, as described below. First, we consider the “ground truth” signals (i.e., not contaminated by cross-talk) *x*_1_(*t*_*i*_) and *x*_2_(*t*_*i*_) for channel 1 and 2, respectively, where *t*_*i*_ (*i* = 1, 2, 3, …) are discrete sampling times. For simplicity, we assume that *x*_1_ and *x*_2_ are time series of independent Gaussian white noise with the same mean *E*[*x*_1_] = *E*[*x*_2_] = 0 and the same variance Var(*x*_1_) = Var(*x*_2_) = *σ*^2^.

**Figure 1.**
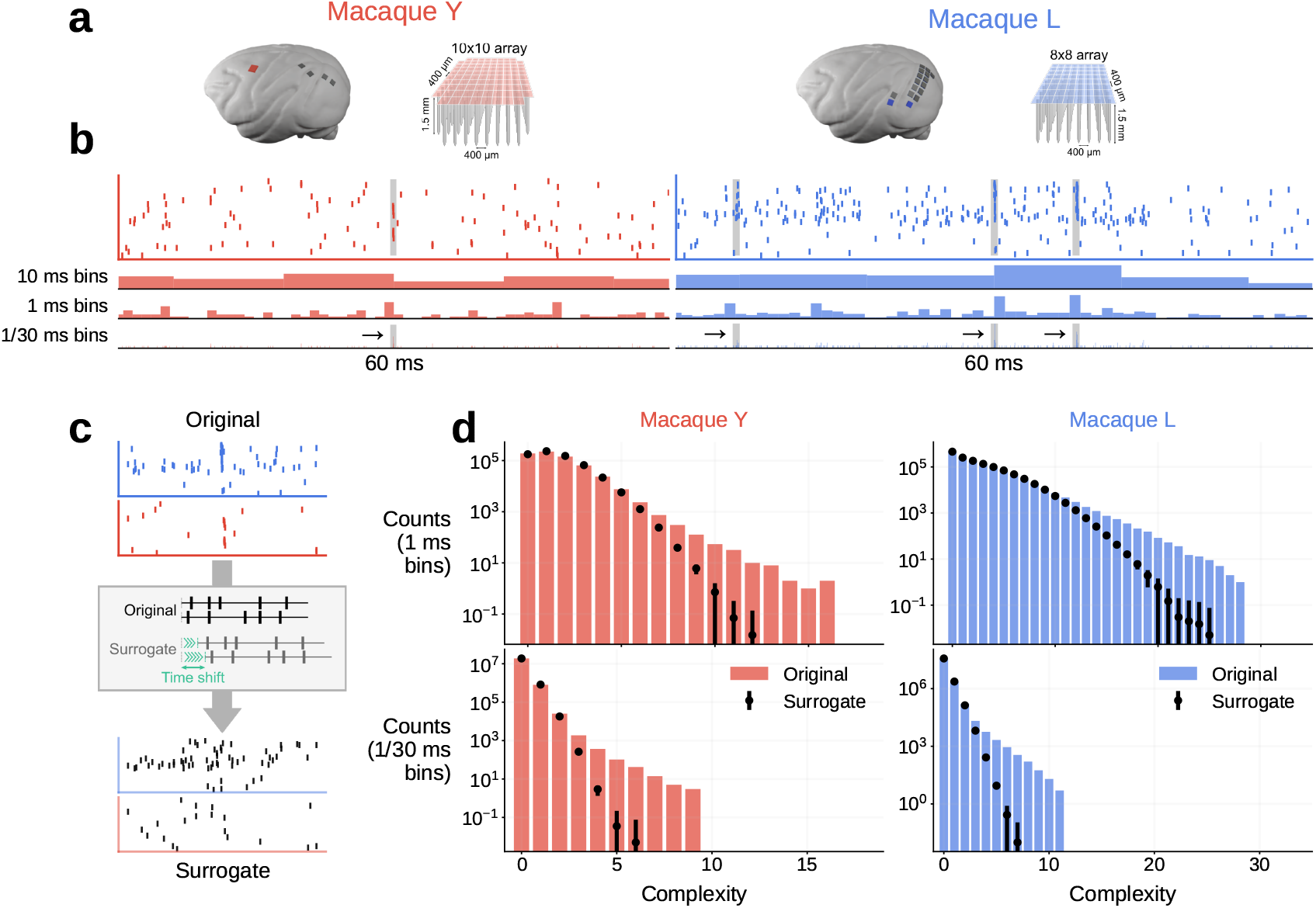
Observation of hyper-synchronous event (HSEs). (a) Electrode arrays and implantation sites for the two macaques, macaque Y: 6x6 electrode arrays in V1, V2, DP, 7A and 10x10 in M1/PMd (we focus here only on the 10x10 array); macaque L: 14 8x8 arrays in V1 and two in V4 (focus here on an array pair with one in V1 and one in V4). The arrays focused on are colored. (b) Raster display (top) and population histograms with different bin sizes (bottom) for a 60 ms long data slice. HSEs are highlighted with gray. (c) Illustration of the surrogate generation method. Original spike trains are shifted against each other by a random amount of time. (d) Complexity distribution of the original data (colored bars), and the mean and standard deviation (dot and error bar, respectively) for the complexity distribution of the respective surrogates. Top: 1 ms bin size; bottom: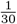 ms bin size.

**Figure 2.**
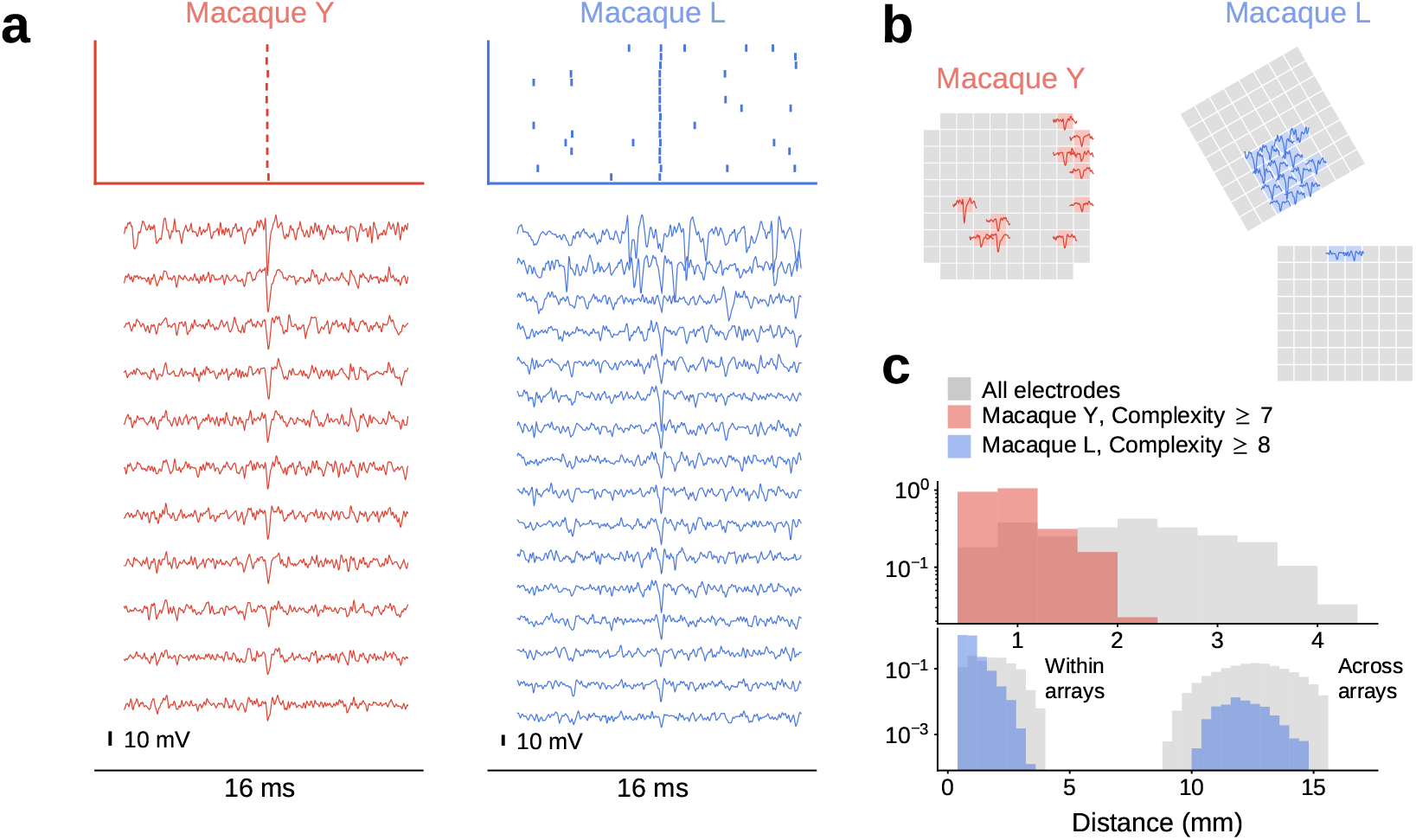
HSEs in relation to the band-passed signals, for macaque Y (red) and L (blue). (a) Dot display (top) centered around a single HSE and the corresponding band-pass filtered (500 Hz - 7500 Hz) raw signals of the channels participating in the HSE. Spike-like excursions are aligned on the HSE. (b) Positions of the electrodes on the respective array for the band-passed signal channels participating in the HSE in (a). The trace of the spike-like excursion in each channel is shown at the corresponding position. (c) Distances between the electrodes involved in all HSEs with high complexity, selected based on the surrogate analysis from Figure 1. For reference, the distance distribution between all possible electrode pairs is shown in gray.

Next, we model the cross-talk as a linear mixing of these two signals. Concretely, we define the measured signals *s*_1_(*t*_*i*_) and *s*_2_(*t*_*i*_) for channel 1 and 2, respectively, as:

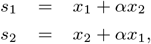

where *α* is a parameter representing the strength of the cross-talk: *s*_1_ and *s*_2_ are identical when *α* = 1, and they are independent when *α* = 0. By this construction, both *s*_1_ and *s*_2_ obey a Gaussian distribution with a mean *E*[*s*_1_] = *E*[*s*_2_] = 0 and variance Var(*s*_1_) = Var(*s*_2_) = (1 + *α*^2^)*σ*^2^. The Pearson correlation coefficient *c*_12_ for the signals *s*_1_ and *s*_2_ is derived as:

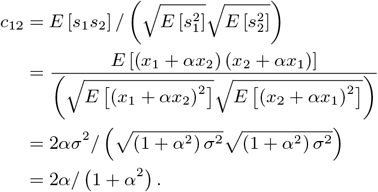

We further introduce spikes to the signals. Here we model spikes in channel 1 by adding a value *A*_1_, representing the spike amplitude, to *x*_1_ at various, random time points. We parameterize *A*_1_ as 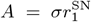, where 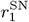 is the signal-to-noise ratio of the spike waveform, such that SUAs with different spike ampli-tudes are represented by modifying the value of 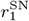. The spikes introduced in the ground truth signal *x*_1_ are transferred to the measured signals *s*_1_ and *s*_2_ via the cross-talk. To count those transferred spikes, we extract spikes from *s*_1_ by thresholding it at 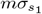, where 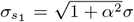 is the standard deviation of *s*_1_ and *m* is a multiplier determining the threshold in relation to the standard deviation. Assuming that *N*_1_ spikes were introduced to *x*_1_, we can derive the expected number *n*_11_ of the spikes transferred from *x*_1_ and detected in *s*_1_ (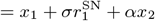 at spike times) as:

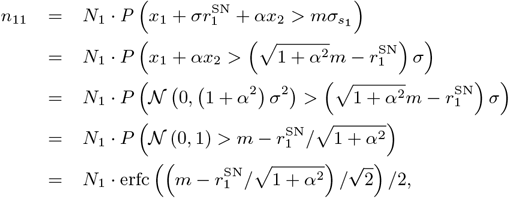

where erfc(·) is the complementary error function.^1^

Note that the factor 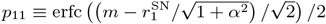 in front of *N*_1_ in the above expression represents the probability that a spike transferred from the ground truth signal *x*_1_ is detected in the measured signal *s*_1_. In a similar manner, the probability *p*_21_ that a spike originally in *x*_1_ is detected in *s*_2_ is derived as 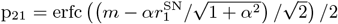. Fur-thermore, assuming *N*^2^ spikes in the other ground truth signal *x*^2^, the probabilities *p*_12_ and *p*_22_ that those spikes are detected in the measured signals *s*_1_and *s*_2_, respectively, are de-rived as 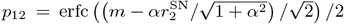 and 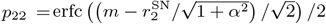

With these probability values, the expected numbers *n*_1_ and *n*_2_ of spikes detected in the measured signals *s*_1_ and *s*_2_, respectively, are represented as:

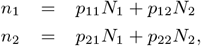

given the numbers *N*_1_ and *N*_2_ of spikes in the ground truth signals *x*_1_ and *x*_2_, respectively. Note that the terms *p*_12_*N*_2_ and *p*_21_*N*_1_ represent the number of spikes transferred from the other channel, and hence these spikes are detected in the both of the measured signals. Thus, the sum of these terms: *n*_1,2_ = *p*_12_*N*_2_ + *p*_21_*N*_1_ represents the number of artifacts in this pair of measured signals. Using this, the HSE index *I*_1,2_ between the measured signals *s*_1_ and *s*_2_ is expressed as:

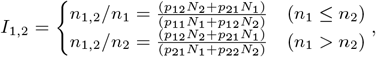

which takes a value between 0 and 1 depending on the parameters.

The model can be extended to represent data with more than a pair of channels. Following the same steps as for the channel-pair model, we first define the ground truth signal *x*_*i*_ for channel *i* (1 *< i < N*_ch_, where *N*_ch_ is the total number of channels) as Gaussian white noise time series with the mean *E* [*x*_*i*_] = 0 and the variance 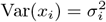. Then, for each channel *i*, spikes are introduced to the ground truth signal by subtracting a value 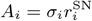, where 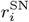 is the signal-to-noise ratio of the spike waveform in channel *i*, at random timings. The measured signal *s*_*i*_ for channel *i* is then obtained as:

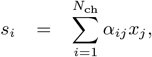

where *α*_*ij*_ is a cross-talk strength matrix, representing the strength of the cross-talk between channels *i* and *j*, taking a value between 0 and 1.

For the simulation of the extended multichannel model shown in Figure 4, we used the following parameters: *N*_ch_ = 100, *σ*_*i*_ = 1 (for all *i*), and 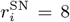 (for all *i*). The cross-talk strength matrix *α*_*ij*_ is defined as follows: i) the diagonal elements *α*_*ii*_ are all set to 1; ii) for the elements in the upper triangle, i.e., *α*_*ij*_ (*i < j*), their values are drawn from a uniform distribution between 0.1 and 0.2 for { (*i, j*) | 0 ≤ *i* ≤ 19, 0 ≤ *j* ≤ 19, *i < j*} ; iii) values for {(*i, j*) | 40 ≤ *i* ≤ 59, 40 ≤ *j* 59, ≤ *i < j* }and { (*i, j*) | 80 ≤ *i* ≤ 99, 80 ≤ *j* ≤ 99, *i < j* }are set in the same manner; iv) values for all the remaining upper triangle elements are drawn from a uniform distribution between 0 and 0.1; and finally, v) values of the elements in the lower triangle are set such that *α*_*ji*_ = *α*_*ij*_ . This results in three groups of 20 channels strongly cross-talking with each other within each group, with weak cross-talks between channels not belonging to the same group. This structure is meant to mimic the correlations between channels observed in the macaque Y data set, as shown in Figure S2. The firing rate the spike trains in the ground truth data is set to 20 Hz for all channels. With this setup, the ground truth signals are generated for a duration of 100 s with the temporal resolution of 0.1 ms. When introducing spikes to the ground truth data, to give a finite temporal width to spikes, the spike amplitude value *A*_*i*_ is subtracted from the background Gaussian white noise time series at 3 successive data points (corresponding to a spike width of 0.3 ms) starting at each spike time.

**Figure 3.**
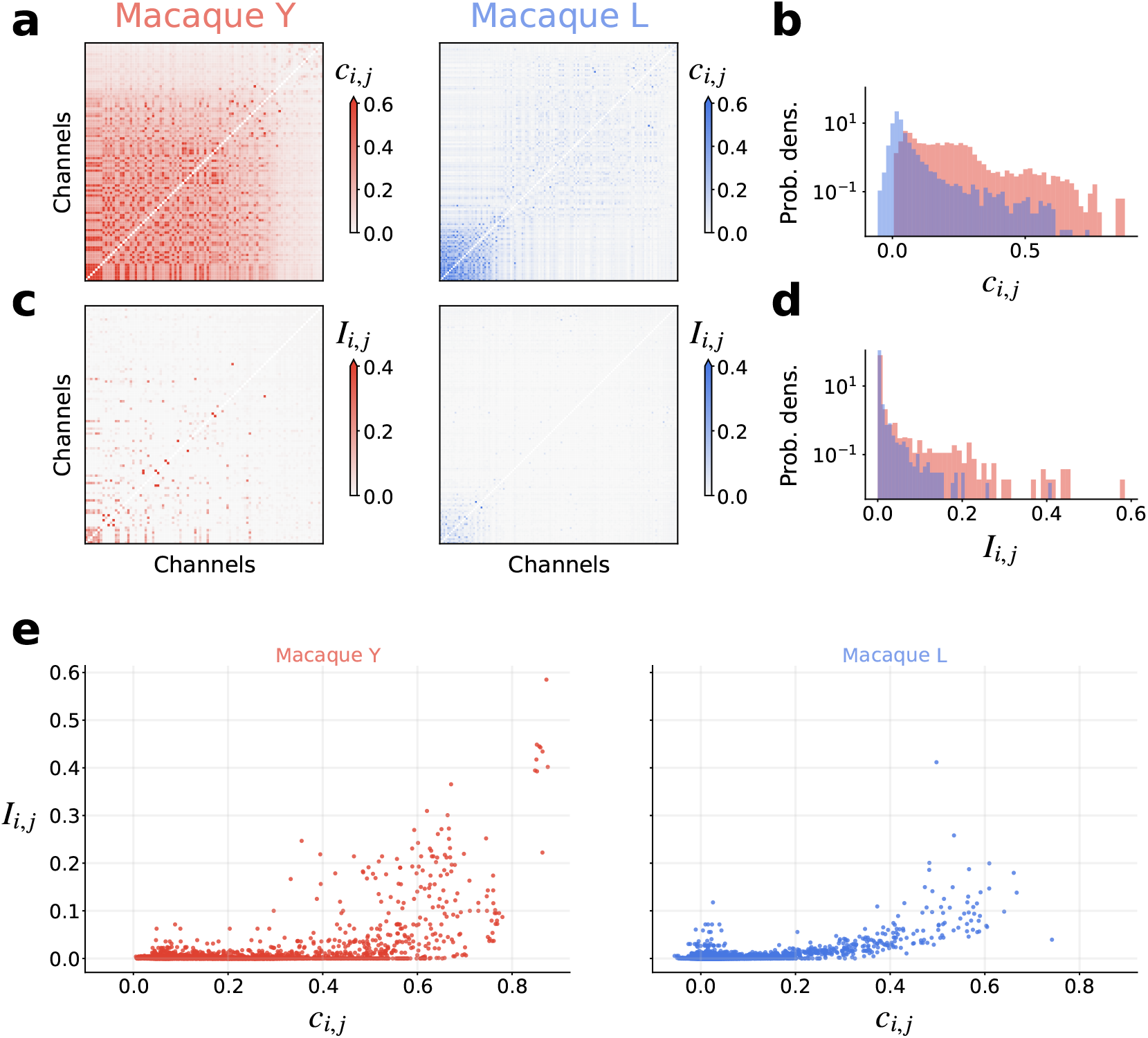
Relation of pairwise band-passed signal correlations to respective pairwise occurrences of HSEs. (a) Pairwise correlation coefficients *c*_*i,j*_ of the band-passed signals. Data are shown in matrix form; the channels are sorted by largest maximum correlation. (b) Distribution of the *c*_*i,j*_ values that appear in the matrix shown in (a). (c) HSE index *I*_*i,j*_ for all channel pairs shown in a channel-by-channel matrix form. The channels are sorted in the same order as in (a). (d) Distribution of the *I*_*i,j*_ values that appear in the matrix shown in (c). (e) Scatter plot of the band-passed signal correlation coefficient *c*_*i,j*_ against HSE index *I*_*i,j*_ of the corresponding channel pair. Each dot represents one channel pair.

**Figure 4.**
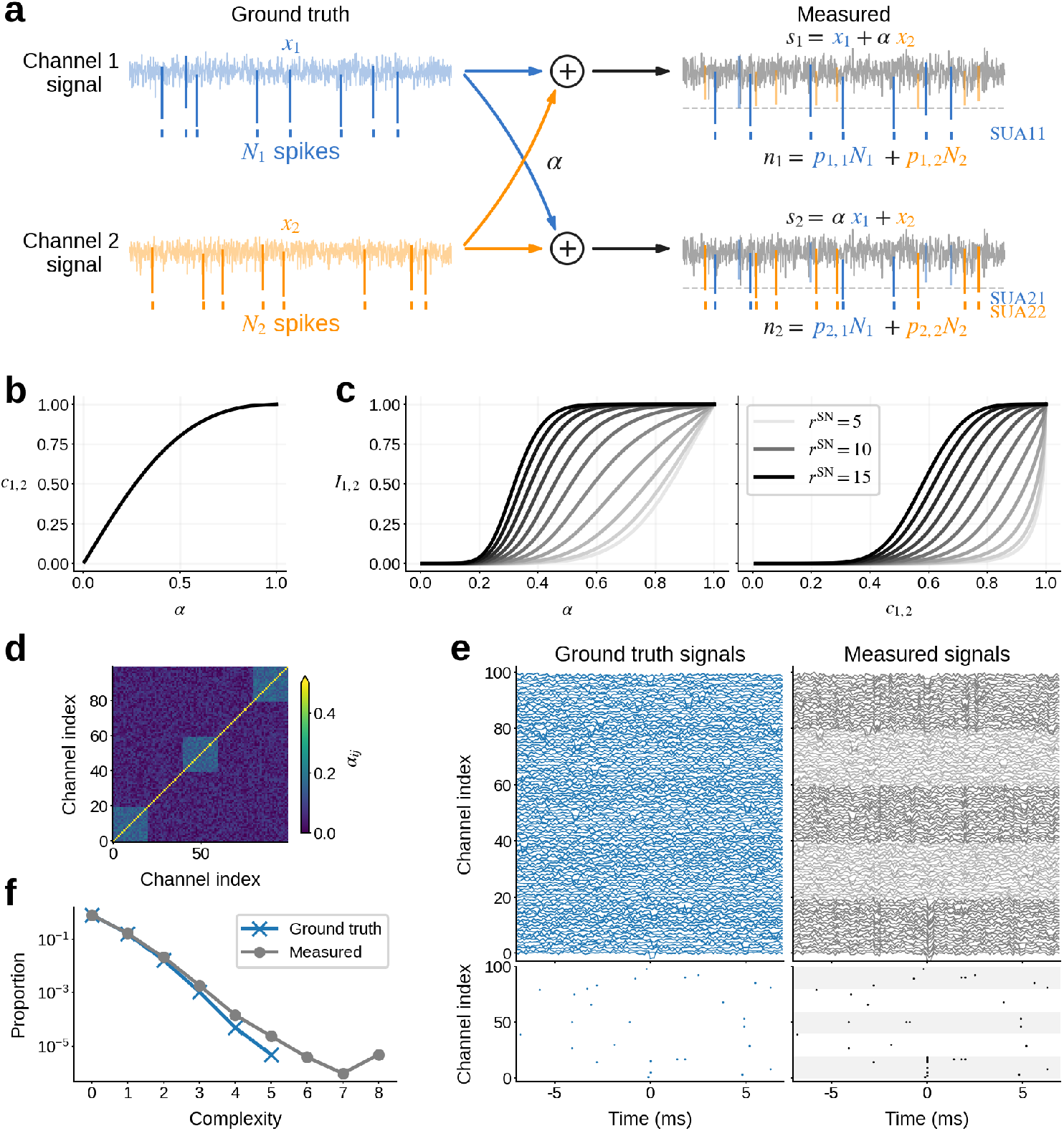
Cross-talk model of artifacts. (a) Illustration of the setup for derivation of the HSE index. Ground truth signals for channels 1 and 2 are mixed, representing the cross-talk, to yield the signals measured in a pair of channels. Spikes (vertical lines) are introduced to the ground truth signals, and they “bleed” into the measured signal of the other channel via the cross-talk. (b) The correlation *c*_1,2_ between the channels plotted against the cross-talk strength *α*. (c) The HSE index *I*_1,2_ of the channels plotted against the cross-talk strength *α* (left) and the correlation *c*_1,2_ between the channels (right). Here the signal-to-noise ratios of spikes in *x*_1_ and *x*_2_ are set to be identical 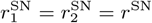, and varied between 5 and 15 in steps of 1, while keeping the threshold multiplier at *m* = 5. The curves for different values of *r*^SN^ are plotted in different shades of gray. (d) The cross-talk strength matrix used for the simulation of the extended multi-channel model shown in panel e. (e) Simulated ground truth (left) and mixed (right) signals of the multi-channel model. The lower panels show the spike trains extracted from the signals in the upper panels. In the plot of the mixed signals, traces for the channels that belong to a strongly cross-talking group are drawn in black, while traces for the channels not participating in the groups are drawn in gray. Accordingly, the background of the spike trains in the lower panel is shaded in gray for the channels that belong to a strongly cross-talking group. (f) The complexity distributions of the extracted spikes of the ground truth (blue) and the mixed (gray) signals computed from the simulations (panel e). The bin width for the complexity calculation is the same as the simulation time step, i.e., 0.1 ms.

### Statistical analysis

*Complexity and estimation of chance levels* To study fine-temporal correlation and detect potential synchronous artifacts in parallel spike train data, we first bin the time axis of the spike trains with a predefined bin width (in the present study we use 1, 0.1, and 1/30 ms bin widths) and compute the complexity of spiking activity at each bin, i.e., the number of units contributing spikes to that bin. Then we make a histogram of the complexity values for all the bins throughout the recording, called complexity distribution (Grün et al., 2008). For easier comparison of the results from different data sets, the histogram is normalized by dividing the counts of all complexities (including the complexity of zero, i.e., bins with no spikes) by the total number of bins, such that the sum of the histogram values over all complexities equals to unity.

We compare the empirical distribution to chance levels from independent data, to elucidate if the real data contains excess spike synchrony. To estimate the complexity distribution of independent data, we use surrogate data, i.e., modified versions of the original data where spike times are intentionally altered. Stella et al. (2022) compared a number of surrogate methods, and according to that, we employ here the ‘time-shift’ method (with a 30 ms shift width, (Pipa et al., 2008)), by which the spike trains are randomly shifted in time against each other. Time shifting destroys potential correlations between spike trains, while conserving many other features of the data such as the inter-spike interval distribution, the firing rate modulations and autocorrelation (Stella et al., 2022). We generate 200 surrogate spike train data sets, and for each of them we compute the complexity distribution. The mean and standard deviation of the count are calculated for each complexity, and plotted together with the empirical complexity distribution for comparison.

*Cross-correlation of band-passed signals* The correlation *c*_*ij*_ between the band-passed signals of channel *i* and *j* is evaluated by the Pearson correlation coefficient as:

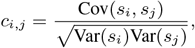

where *s*_*i*_ and *s*_*j*_ represent the band-passed signals of channel *i* and *j*, respectively, obtained as described in Signal processing, and Var(*s*_*i*_) and Cov(*s*_*i*_, *s*_*j*_ ) represent the variance of *s*_*i*_ and the covariance between *s*_*i*_ and *s*_*j*_, respectively.

*Measure of spike synchrony* Assume that we have *n*_*i*_ spikes in channel *i* at times 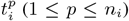. We discretize the time axis in bins of width *b*, which should be small enough to contain at most one spike in a bin, and obtain a set of bin indices 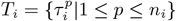, which indicates the positions of the time bins where the spikes are observed (in other words, the time of the *p*-th spike is in the time range 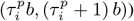. The number *n*_*i,j*_ of spike synchrony events between channels *i* and *j* is obtained as *n*_*i,j*_ = |*T*_*i*_ ∩ *T*_*j*_ .| We define the pairwise HSE index *I*_*i,j*_ as

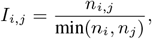

which is a measure of the abundance of spike synchrony events between a pair of channels *i* and *j*. We choose to divide by the smaller spike count so that the HSE index is independent of the channel order (i.e., *I*_*i,j*_ = *I*_*j,i*_) and also sensitive to HSEs occurring on channels with small number of spikes.

In a similar manner, we define the global HSE index *I*_*i*_ as

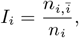

where 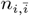 is defined as 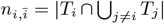, which represents the number of spike synchrony events between channel *i* and any other channel. Thus, the global HSE is a measure of the abundance of spike synchrony events between channel *i* and any other channel.

## Results

### Synchronous spike events are ubiquitous in multi-electrode systems

To demonstrate how artifacts appear in massively parallel spike train data, we start inspecting two macaque data sets from different laboratories and animals (Figure 1a). In the raster-display of the simultaneously recorded spike trains (Figure 1b, top), we notice events where many neurons exhibit spikes simultaneously (highlighted in gray). To examine the temporal precision of these spike coincidences, we compute the population spike time histograms with different bin sizes (Figure 1b, bottom). At the resolutions of 1 ms and 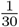 ms, we find some bins with outstandingly high spike counts, not visible at the resolution of 10 ms. We term these extremely precise spike coincidences within 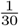 ms as hypersynchronous events (HSEs).

We further quantify the HSEs by their complexity (i.e., number of involved neurons) (Grün et al., 2008) and compare the obtained complexities to those expected by chance using a surrogate method (Stella et al., 2022) (Figure 1c; see Materials and Methods for details). Figure 1d shows the distributions of complexity values from the original and the surrogate data, for two different bin sizes (1ms and 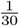 ms). For both bin sizes, the original data contains more high complexity HSEs than expected by random chance. Considering that the co-ordination of spike times on the timescale of 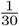 ms is very unlikely, we conclude that many of the HSEs in the experimental data are artifacts, which we name synchrofacts.

We find artifacts in two separate data sets originating from two different laboratories, recorded in two different cortical areas of different macaques. Therefore we suspect that artifacts are common across recordings of the same type. While both data sets were recorded with Blackrock recording systems, artifacts also occur in other recording systems, e.g. in Neuropixels recordings (see section High density probe recordings also contain artifacts). Taken together, our observations highlight the ubiquity of artifacts.

One might think that the data can be cleaned by simply removing all HSEs. However, that is not an appropriate solution to get rid of artifacts, since this also removes many genuine neuronal spikes and, thus, genuine neuronal synchrony. One can demonstrate this by removing all spikes in the bins with complexity of two or larger at 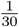 ms resolution, and then calculating the complexity distribution, to be compared to the complexity distribution of its surrogates. The surrogate distribution obtained in this way contains complexities of two or higher that occur by chance. However, the original distribution after the removal totally lacks complexities higher than 2, and hence it differs vastly from the surrogate distribution (see Supplementary Figure S1). Thus, removing all HSEs significantly alters the statistical properties of the spike trains by selectively eliminating higher-order chance synchrony, which can lead to false results regarding fine temporal correlations in the spike trains.

To summarize, HSEs in massively parallel spike trains commonly contain synchrofacts, which are artificial spike synchrony events on a timescale of the data sampling frequency. Synchrofacts can only be spotted when analyzing the data on very fine timescale and show themselves as excess HSEs. Since HSEs contain not only artifacts but also genuine neuronal synchrony (as well as chance coincidences), removing all HSEs for cleaning the data is not a proper approach. Hence, we aim at discriminating artifacts from chance coincidences of real spikes, for selectively removing artifacts from the data.

### Synchronous spike events with nearly identical waveforms are not of neural origin

Following the observations in the previous section, we now take a deeper look into the HSEs for features discriminating artifacts from chance spike coincidences. Whereas an HSE with a very high complexity is likely to be an artifact, we cannot rule out the possibility that it is a real spike synchrony. In fact, there is no way of discriminating between them on the description level of spikes. Thus, seeking for additional information to spot artifacts, we inspect the raw signals from which the spike events are extracted.

Here for the macaque data sets, the spike extraction was performed by thresholding the band-pass filtered raw signals in the frequency range of 500 Hz − 7500 Hz (see Signal processing in Materials and Methods for details). For simplicity, we henceforth refer to the band-pass filtered raw signal merely as band-passed signal. Figure 2a shows the band-passed signals of multiple channels around the timing of one HSE. We observe highly similar spike-like waveforms in multiple channels simultaneously, but with different amplitudes. Furthermore, the ongoing signal fluctuations before and after the HSE appear to be correlated across these channels. The electrodes of these channels are not necessarily in close spatial proximity, as shown in Figure 2b. To quantify this observation, we measure the distances between the electrodes of channels participating in each and every HSEs with high complexities. The distribution of the obtained distances (Figure 2c) shows that the high complexity HSEs often involve electrodes that are far apart, up to several millimeters. In particular, in the case of macaque L, HSEs appear even across the two arrays located on opposite banks of the lunate sulcus.

To summarize, we have observed i) sub-millisecond synchronization of spikes across channels, ii) nearly identical waveforms of those spikes, iii) visible correlation between band-passed signals, and iv) partly large distances between the participating electrodes. Taken together, these observations strongly suggest that many of these events are artifacts due to electric cross-talk between channels, rather than multiple electrodes recording the same neuron or some novel form of fast neuronal synchronization.

### Cross-correlation and hyper-synchronous events can detect artifacts

An electric cross-talk between a channel pair is expected to increase the Pearson correlation coefficient between the band-passed signals of these channels. Figure 3a shows the correlation coefficient *c*_*i,j*_ between the band-passed signals of channels *i* and *j* (see Cross-correlation of band-passed signals in Materials and Methods for details), for all channel pairs in the two data sets in a matrix form (only for *i* ≠ *j*). The distribution of the *c*_*i,j*_ values in the matrix is shown in Figure 3b. The coefficients are mostly larger than zero, i.e., the band-passed signals are generally positively correlated. The coefficient values vary strongly across channel pairs, with some pairs showing quite strong correlation, in both recording setups from different laboratories.

There may be multiple possible origins for such strong correlations, ranging from damages on the electrodes to electric shortcuts between channels due to various reasons, e.g., crossing of the implanted cables below the skull, dirt in the headstage connector, interference between the cables connected to the amplifier, etc. (Yu et al., 2009). As shown in Figure 2b and c, the artifacts are not limited to spatially proximal channels but extend over the whole array or even across arrays, indicating that a deficit in a local group of channels cannot be the primary reason for these correlations. Another possibility is a local deficit on the headstage connector, but this is also denied by examining the spatial extent of correlated channels mapped on the connector (see Macaque L data in Supplementary Figure S2). Thus, no single reason can explain the observed correlations, but rather there would be multiple causes residing on different stages of the setup. Eliminating all possible causes from the setup is practically not feasible, and hence we cannot avoid having such correlations in the data.

For channel pairs with a large *c*_*i,j*_ value, their bandpassed signals must be similar to each other. Hence, if one channel of such a pair had spike-like waveforms, the other channel should also have those at the same time. This leads to the expectation that a channel pair with a larger *c*_*i,j*_ should show more artifacts. To confirm this, we first count, for each channel pair, the number of HSEs containing spikes from that pair. We then define the HSE index *I*_*i,j*_ of channel *i* and *j* as the ratio of the obtained HSE count to the spike counts of these two channels (see Measure of spike synchrony in Materials and Methods for the formula). Figure 3c shows the *I*_*i,j*_ values obtained from the two data sets in a matrix form (only for *i* ≠ *j*). The distribution of those values in the matrixis shown in Figure 3d. We find *I*_*i,j*_ values ranging from zero to 0.5, while the value expected from an independent pair of spike trains is on the order of 10^−3^ (see Supplementary Figure S3).

We then plot *I*_*i,j*_ against the correlation coefficient *c*_*ij*_ to look for a systematic relation between them (Figure 3e). We find that a majority of channel pairs have HSE index values close to zero, meaning that hardly any spikes are shared by these pairs. Especially the channel pairs with low correlation coefficients *c*_*i,j*_, such as *<* 0.4, have very low HSE index values. As the correlation *c*_*i,j*_ gets larger than 0.4, the HSE index value rapidly increases, indicating a critical correlation beyond which spikes becomes very likely to be detected in both channels. Note that an HSE index larger than zero does not immediately indicate artifacts. As shown in Figure 1d, a certain amount of HSEs can be explained by chance. However, as we mentioned before, the HSE index expected from independent spike trains is very small, on the order of 10^−3^. Hence, an HSE index excessively greater than this strongly indicates presence of artifacts.

### Cross-talk model explains the emergence of artifacts

To understand the origin of the relation between the band-passed signal correlation *c*_*i,j*_ and the HSE index *I*_*i,j*_, here we present a simple model of cross-talk between channels (see Cross-talk model in Materials and Methods for details). This model enables us to derive an analytic expression of the relation, which is in agreement with our experimental observations (Figure 3e).

Our model assumes two Gaussian white noise (mean 0, variance *σ*^2^) time series *x*_1_ and *x*_2_ as the cross-talk-free “ground truth” signals. We model the cross-talk as a linear mixing of these two signals (Figure 4a), such that the mixed signals *s*_1_ and *s*_2_ measured on channel 1 and 2, respectively, are:

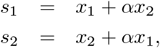

where 0 ≤ *α* ≤ 1 is the strength of the cross-talk: *s*_1_ and *s*_2_ are identical with *α* = 1, and independent with *α* = 0. The Pearson correlation coefficient *c*_1,2_ between *s*_1_ and *s*_2_ can be written in terms of *α* as (see Cross-talk model in Materials and Methods for derivation):

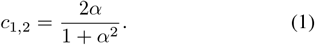

Thus the correlation *c*_1,2_ increases nonlinearly with the crosstalk strength *α*, as shown in Figure 4b.

For the derivation of the HSE index, we need to introduce spikes to the signals. We model spikes by subtracting a value *A*_*i*_, representing the spike amplitude, from the ground truth signal *x*_*i*_ (*i* ∈ {1, 2 }) at random time points. We parameterize *A*_*i*_ as 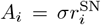, where 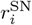 is the signal-to-noise ratio of the spike waveform in *x*_*i*_. Spikes are then extracted from the mixed signals *s*_1_ and *s*_2_ by thresholding, as commonly done for experimental data. We set the threshold for *s*_*i*_ at 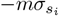, where 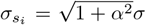 is the standard devia-tion of *s*_*i*_ and *m* is a multiplier (Quiroga et al., 2004). The expected numbers *n*_1_ and *n*_2_ of spikes detected in *s*_1_ and *s*_2_, respectively, are:

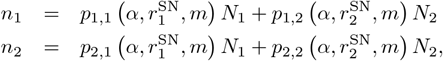

where *N*_1_ and *N*_2_ are the number of spikes introduced in *x*_1_ and *x*_2_, respectively, and *p*_*i,j*_ is the probabilities that a spike from *x*_*i*_ is detected in *s*_*j*_ as a function of the parameters *α*, 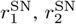, and *m* (see Cross-talk model in Materials and Methods for more details).

The spikes transferred across channels by the cross-talk are detected in both channels, so the sum of the numbers of such spikes, i.e., *n*_1,2_ = *p*_1,2_*N*_2_ +*p*_2,1_*N*_1_ represents the number of artifacts in this pair of channels. We can now calculate the HSE index as *I*_1,2_ = *n*_1,2_*/*min(*n*_1_, *n*_2_) (see Cross-talk model in Materials and Methods for detailed derivations). While this model could be extended to include more realistic details such as different types of noise, spike waveforms, and so on, the simple model as described here suffices for the current purpose of deriving the HSE index analytically.

Through the dependence of *p*_*i,j*_ on the cross-talk strength *α*, the HSE index *I*_1,2_ is dependent on *α* and thereby the correlation coefficient *c*_1,2_ (Figure 4c). Interestingly, hardly any artifacts are observed in the model for *α <* 0.2 or *c*_1,2_ *<* 0.4, suggesting that some degree of cross-talk might be tolerable in recording systems. However, for higher *α* or *c*_1,2_, the HSE index rapidly increases in the form of a sigmoid, depending on the signal-to-noise ratio *r*^SN^. This is in good agreement with what we observed in our macaque datasets (Figure 3e).

In the real data, the signal of one channel is typically correlated not only with another channel but with multiple other channels at once, as seen in Figure 3. The model can be extended to represent such data by introducing multiple channels (let *N*_ch_ be the number of channels) and expanding the cross-talk strength parameter *α* to a cross-talk strength matrix *α*_*ij*_ (1 ≤ *i* ≤ *N*_ch_, 1 ≤ *j* ≤ *N*_ch_), which represents the cross-talk strength between channel *i* and *j* (see Cross-talk model in Materials and Methods for more details).

Here we demonstrate that the extended multi-channel model can simulate artifacts and correlated signal fluctuations as seen in our experimental data (Figure 2a), given a proper cross-talk strength matrix. We consider a model of 100 channels and a cross-talk strength matrix as shown in Figure 4d. The diagonal elements of the matrix are all set to 1 (to be consistent with the channel-pair model shown above), and the remaining elements are defined such that the data contains three groups of 20 channels (i.e., channels 0-19, channels 40-59, and channels 80-99) with strong cross-talk within each group. The ground truth signal of each of the 100 channels is generated independently as a Gaussian white noise time series with a Poisson spike train (20 Hz firing rate, 0.3 ms spike width) injected (see Cross-talk model in Materials and Methods for more details of the simulation), as shown in Figure 4e, left. Since the injected spike trains are independent between channels, the complexity distribution of the spikes in the ground truth signals is just as expected from chance spike coincidences between channels (Figure 4f, blue). From these ground truth signals, mixed signals are generated according to the cross-talk strength matrix (Figure 4e, right). Strong correlations in the signals among the channels within each cross-talking channel group (black traces) are evident, compared to the correlations among the channels not participating in those groups (gray traces). As a consequence, when there is a chance spike coincidence in the ground truth signals of a cross-talking channel group (e.g., at the middle of Figure 4e, left, for the bottom channels), the strong cross-talk within the group replicates those spikes into the other channels of the group at the same timing, resulting in a formation of a artifact (Figure 4e, right, for the bottom channels). Accordingly, the complexity distribution of the mixed signals exhibits a heavy tail of high order complexities (Figure 4f, gray), which resembles the complexity distribution of the real data (Figure 1d).

### ZCA whitening effectively removes artifacts

From this model we now see that the correlations between the signals lead to emergence of artifacts. Conversely, the model poses a possibility that if one could properly decorrelate the measured signals, the ground truth signals could be recovered and the artifacts would be dissolved. Since our model assumes a linear mixture of the ground truth signals via symmetric cross-talks, the measured signals can be decorrelated by whitening them based on their covariance matrix, a method called zero-phase component analysis (ZCA) whitening (Bell and Sejnowski, 1997; Kessy et al., 2018).

In Figure 5a we illustrate an example application of ZCA whitening to simulated data of the model. Similarly to the example shown in the previous figure (Figure 4e), a chance spike coincidence in the ground truth signal develops to a artifact in the mixed signal due to cross-talk (Figure 5a, left and center). After an application of ZCA whitening to the mixed data, the ground truth signals are perfectly recovered for all channels (Figure 5a, right), and the spikes that compose the artifact are removed except for the two that are the original chance coincidence spikes that caused the artifact. All the other spikes outside the artifact are also kept intact. In fact, ZCA whitening is exactly the inverse operation of the linear mixture in the proposed cross-talk model. Thus, ZCA whitening performs perfectly to recover the ground truth signals as long as the correlations in the mixed signals are generated according to the cross-talk model.

**Figure 5.**
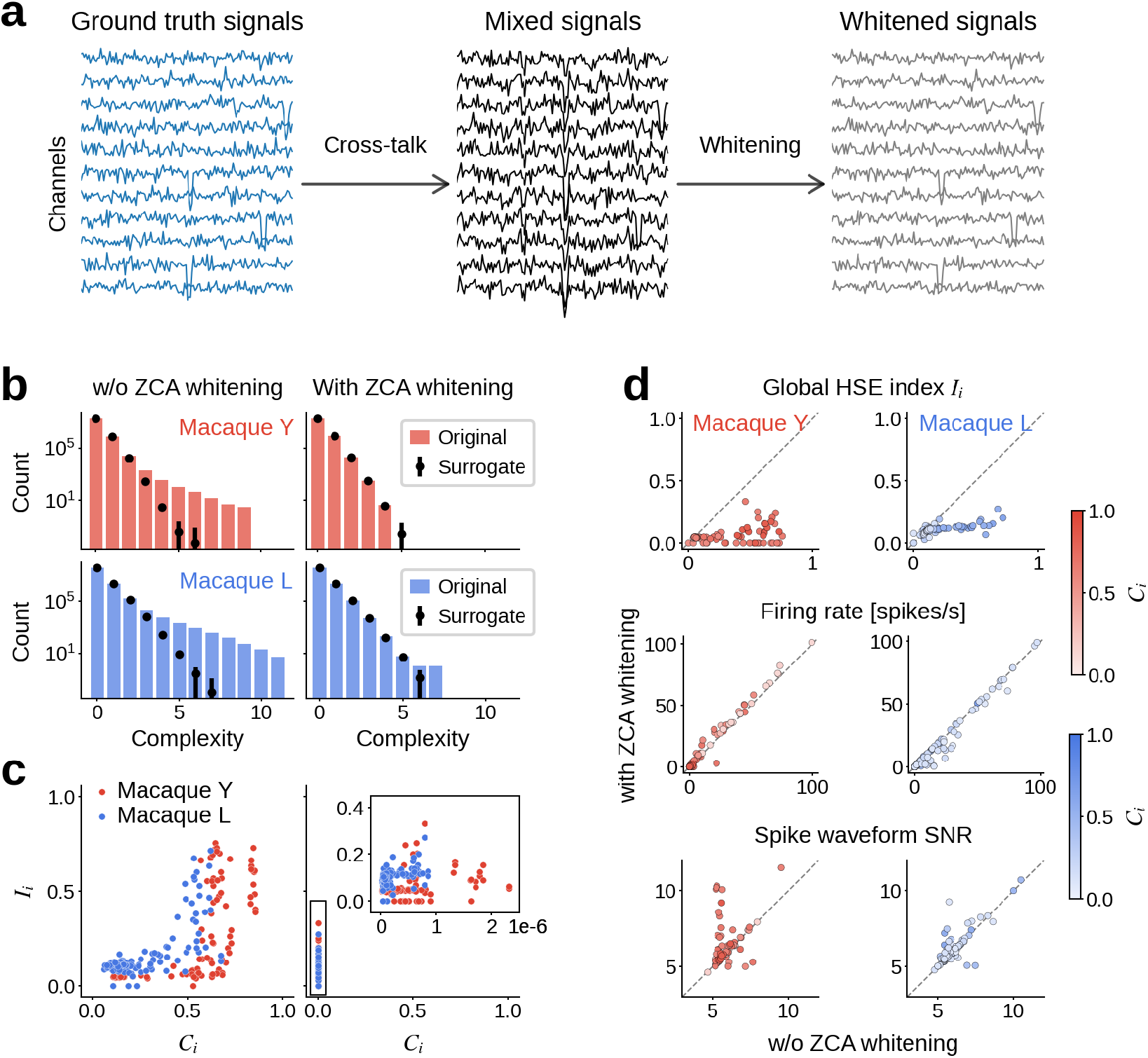
Removal of artifacts by use of ZCA whitening. (a) Example application of ZCA whitening to simulated data of the cross-talk model. ZCA whitening applied to the mixed signals from the model (center) yields whitened signals (right), which are identical to the ground truth signals (left) used to generate the mixed signals. (b) Complexity distribution of the Macaque Y (red bars) and Macaque L (blue bars) data, without (right) or with (left) ZCA whitening. For comparison, the mean (black dot) and standard deviation (error bar) of the complexity distribution of respective surrogate data are also shown. (c) Global HSE index *I*_*i*_ for channels of the Macaque Y (red) and Macaque L (blue) data, plotted against the highest correlation *C*_*i*_ of the channels, without (left) or with (right) ZCA whitening. The inset in the right panel shows a magnified view of the same data. (d) Comparison of spike train statistics of the Macaque Y (left) and Macaque L (right) data between without (x-axis) and without (y-axis) ZCA whitening. The color of dots represents the highest correlation of the respective channels without ZCA whitening, as indicated in the color bars to the right. Top: global HSE index for each channel. Middle: firing rate of spikes in each channel. Bottom: signal-to-noise ratio (SNR) of spike waveforms in each channel.

Now we apply ZCA whitening to our experimental data and see if this method can remove the artifacts. Figure 5b shows the complexity distributions for the macaque Y (top right) and macaque L (bottom right) data with application of ZCA whitening (and for comparison, the corresponding distributions without ZCA whitening (Figure 5b, left), which are the same data as in Figure 1d). With ZCA whitening, the complexity distribution matches quite tightly to the distribution from surrogate data, i.e., independent data, with a considerable reduction of high order complexities seen in the complexity distribution without ZCA whitening.

To quantify the effects of ZCA whitening on single channels, we introduce a measure evaluating the participation of each channel in HSEs. We term this as the global HSE index *I*_*i*_ for channel *i*, defined as the ratio of the number of HSEs including channel *i* to the number of all spikes on channel *i* (see Measure of spike synchrony in Materials and Methods for the formula). The HSE index *I*_*i,j*_ that we have introduced before for a pair of channels *i* and *j* is henceforth referred to as pairwise HSE index. Furthermore, we also introduce a channel-wise measure of signal correlations, i.e., the highest correlation coefficient for each channel i to all other channels, denoted as *C*_*i*_ for channel *i* and defined as *C*_*i*_ = max_*j*≠*i*_(*c*_*i,j*_), where *c*_*i,j*_ denotes the correlation coefficient between the band-passed signals of channel *i* and *j*. Figure 5c shows, for data with (right) or without (left) ZCA whitening, the global HSE indices *I*_*i*_ plotted against their respective highest correlation coefficients *C*_*i*_ for all channels in macaque Y and L data sets. In the case without ZCA whitening (Figure 5c, left), as expected from the model, the channels with high *C*_*i*_ values are indeed also the channels with high *I*_*i*_ values, while the channels with *C*_*i*_ below 0.4 show rather small *I*_*i*_. On the other hand, in the case with ZCA whitening (Figure 5c, right), all channels show almost zero correlations, which is a direct consequence of the whitening, and also show considerably lower *I*_*i*_ values, indicating an overall suppression of HSEs by ZCA whitening.

Figure 5d shows the statistics of the two data sets with (y-axes) and without (x-axes) ZCA whitening. Figure 5d (top) compares the global HSE index *I*_*i*_ with and without ZCA whitening. All channels show a reduction in *I*_*i*_ with ZCA whitening. This reduction is achieved without reduction in the firing rates of the spike trains in a majority of channels, as evidenced by the comparison of channel-wise firing rate with and without ZCA whitening (Figure 5d, mid-dle). Most channels show similar firing rates with and without ZCA whitening, indicating that ZCA whitening removes only a tiny fraction of spikes, which are most likely the ones copied by the cross-talk between channels. Unexpectedly, in the Macaque Y data set (left column) there are a considerable number of channels which rather increase their firing rates with ZCA whitening, which seems contradictory to the removal of copied spikes. This increase in firing rates is caused by a larger signal-to-noise ratio (SNR) of the spike waveforms, which is due to the reduction of the magnitude of the background fluctuations by the decorrelation of signals. Figure 5d (bottom) shows a comparison of the spike waveform SNR with and without ZCA whitening. For a majority of channels the SNR is increased by ZCA whitening. Thus, in addition to removing spikes copied by cross-talk, ZCA whitening also improves signal quality of single channels through a reduction of the background fluctuations by decorrelation, which makes spike detection more reliable.

### High density probe recordings also contain artifacts

So far we have focused on the occurrence and removal of artifacts in recordings with Utah electrode arrays, where the inter-electrode distance is at least 400 *µm*. Nowadays more and more laboratories start to use high-density electrodes, such as Neuropixels probes (Jun et al., 2017), with interelectrode distances as small as ∼ 20 *µm* at minimum. With such a high electrode density, a spike of one neuron can be recorded at multiple neighboring electrodes, which causes an apparent synchronous spike event among the respective channels. Such apparent spike synchrony is typically sorted into a single unit by use of a waveform template considering signals from multiple neighboring channels, as is done in, e.g., the Kilosort sorting algorithm (Pachitariu et al., 2024). However, this does not necessarily mean that spike trains sorted in such a way are free from artifacts. They can still contain artifacts due to cross-talks between distant channels pairs, or any other causes that introduce strong correlations between signals at distant channels. In this section we focus on spike train data from a Neuropixels recording sorted by Kilosort to show that these data also contain artifacts and our ZCA-whitening-based approach presented above can be used to remove them.

The data we consider here were recorded with a Neuropixels 1.0 probe (see Materials and Methods) in an awake mouse during visual stimulation. The probe was inserted through the primary visual cortex (V1; Figure 6a), with the tip of the probe reaching the superior colliculus (SC). The band-pass filtered raw signals appear largely similar across all channels (Figure 6b, left). This is primarily because all recorded channels are relative to the external reference electrode (in this recording placed on the dura above the cerebellum) and are therefore equally affected by signal fluctuations at the reference. Consequently, the signals can show very strong correlations between channels, even at distant channel pairs (Figure 6c, top).

**Figure 6.**
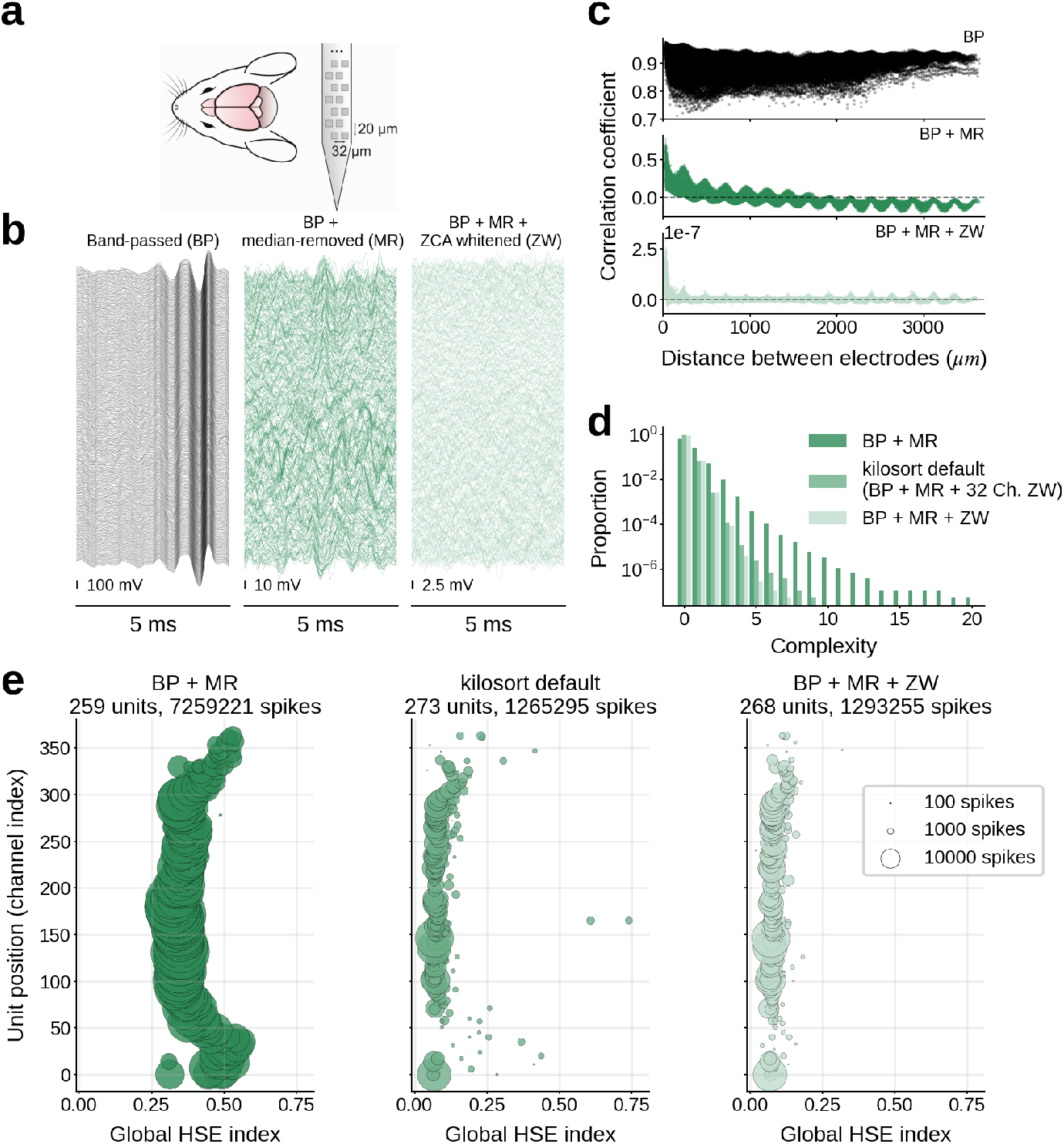
Artifacts in Neuropixels recording. (a) Illustration of the subject animal and the probe. (b) Traces of the recorded signals after progressive signal processing steps. Channels are ordered according to their spatial position on the probe (the bottom corresponds to the tip of the probe). Time and voltage scales are indicated by the respective scale bars. Left: signals after band-pass filtering (BP) in 500-7500 Hz. Center: signals after BP and median removal (MR). Right: signals after BP, MR, and ZCA whitening across all channels (ZW). (c) Pearson correlation coefficient (CC) between the signals of a channel pair as a function of the channel distance. Note the different value ranges on the y-axis for different panels. Top: CC for signals after BP. Middle: CC for signals after BP and MR. Bottom: CC for signals after BP, MR and ZW. (d) Complexity distributions for signals with different signal processing steps. For ‘kilosort default’, BP and MR are applied to the signals, and then ZCA whitening is applied on each neighboring 32 channel groups, as done by Kilosort sorter with its default settings. (e) Global HSE index of sorted single units plotted against their position on the probe. The size of the disk is proportional to the number of spikes of the corresponding single unit, as indicated by the legend in the rightmost panel. The total number of detected single units and spikes are shown in the title of each panel. The dataset labels and the color convention are common to those in panel d.

A standard signal pre-processing step is therefore to first remove signals from electrodes outside of the brain and then to calculate the median of signals across all remaining channels at each time point. By subtracting this median signal from each channel - commonly known as median removal or common average referencing - common noise patterns that are shared across all channels are effectively removed (for more details see Materials and Methods). The signals after this median removal show more variability across channels (Figure 6b, center) and generally contain much less correlation between channels (Figure 6c, middle). However, there are still strong correlations in channel pairs with very short distances, as expected from the conductivity of the brain tissue. Besides that, there are clear negative correlations between channels separated by more than 2000 *µm*, which can be recognized also in the signal traces in Figure 6a (right) for top and bottom channels showing inverted waveforms. These negative correlations seem not to be of physiological origin, because the bottom electrodes are in another brain region than the top electrodes (namely, SC and V1, respectively). One may additionally notice oscillatory modulations of the correlation coefficient as a function of the inter-electrode distance. In the scope of the present study, we do not discuss these oscillations further, but we suspect that they most likely stem from the hardware or get introduced by the correction for the time shifts introduced by the multiplexing during the recording (see Materials and Methods). The time-shift correction is a necessary pre-processing procedure to obtain properly time-aligned signals from the raw recordings.

ZCA whitening on the median-removed signals almost totally gets rid of the positive and negative correlations (Figure 6b, bottom). Kilosort sorting applied to the median-removed and ZCA whitened signals results in a considerable difference in terms of the complexity distribution of the obtained spike trains (Figure 6d): the distribution for the median-removed signal (“BP + MR” in Figure 6d) shows a heavy tail of high order complexities up to 20, while the distribution for the ZCA whitened signal (“BP + MR + ZW” in Figure 6d) shows a faster decay with a maximum complexity of 7.

We need to note that ZCA whitening is already included in the typical and recommended workflow of the Kilosort sorting algorithm as a signal pre-processing step, which we intentionally disabled when we compute the complexity distribution for the median-removed signal in Figure 6c. The ZCA whitening in Kilosort is by default applied to the 32 channels closest to the target channel. In this manner, the signals in the channels separated by more than 32 channels are not decorrelated to each other, i.e., the negative correlations between distant channel pairs we found in the median-removed signals are not removed. This could lead to artifacts between sorted single units in the top and the bottom channels, and in fact, the complexity distribution for the median-removed signals sorted with the Kilosort default settings (“kilosort default” in Figure 6d) shows more high order complexities than our ZCA whitened signals, where ZCA was applied to all channels simultaneously (“BP + MR + ZW” in Figure 6d). To further examine the degree and location of artifacts in these data, we compute the global HSE index *I*_*i*_ for each sorted single unit and plot it against the position of that unit on the probe (Figure 6e). In the case without ZCA whitening (Figure 6e, left), the units around the top and the bottom of the probe stand out with particularly high *I*_*i*_, as we expected from the negative correlations between top and bottom channels. These correlations cause Kilosort to extract spikes in these channels at the same time point just with opposite signs. In the case with ZCA whitening with Kilosort default settings (Figure 6e, center), much less number of spikes are detected and hence units naturally show lower *I*_*i*_ values, but the units around the top and the bottom of the probe still exhibit relatively high *I*_*i*_ values compared to the other units. This is a reflection of the negative correlations that still remain in the signals by the spatially restricted application of ZCA (32 channels in parallel) by Kilosort. In the case with ZCA whitening applied to all channels altogether (Figure 6e, right), a similar number of spikes and units are detected as in the previous case, whereas the units around the top and the bottom of the probe do not show particularly high *I*_*i*_ values, indicating that artifacts including these units are removed. With these results, we conclude that, although the internal cause of artifacts is probably different from the case using Utah array(s), Neuropixel recordings can contain artifacts, and ZCA whitening applied to all channels together effectively removes those artifacts from the data.

## Discussion

We have shown, based on examination of electrophysiological recordings from multiple laboratories, that HSEs in massively parallel spike trains commonly contain artifacts, i.e., artificial extremely precise (at the data sampling resolution) spike synchrony events. These artifacts express themselves in the raw signal as nearly identical activity traces, observed also between distant electrodes with submillisecond precision, strongly suggesting that they originate from external noise or cross-talk in the recording hardware. We introduced a measure, the HSE index (*I*_*i,j*_), to quantify the occurrences of HSEs in each channel pair, and showed its systematic relation to the correlation coefficient (*c*_*i,j*_) of the band-pass filtered raw signals. We also presented a minimal model of cross-talk between signals that explains the observed relation between *I*_*i,j*_ and *c*_*i,j*_, suggesting that cross-talk is the main source for the artifacts in the examined datasets recorded by the Utah arrays. Based on the observations and the model, we suggested cleaning of the data by decorrelating the signals using ZCA whitening, which removes nearly all above-chance HSEs. We in addition showed that recordings with high density probes such as Neuropixels probes may include artifacts and that the ZCA whitening based cleaning method removes these artifacts in those recordings, even though the artifacts there seem not to stem from electric cross-talks between channels. We thus highlight the importance of removing these artifacts from neural data, since they could bias analyses and produce misleading, as will be discussed below. A crucial thing to note is that we cannot distinguish which spikes are artifact spikes and which are “real” spikes. We conclude that these are artifacts in the data from the abundance of HSEs and the resulting high correlation of the spiking activity in the data but do not identify artifacts on a single spike level (with a few exceptions see Figure S4). While spiking activity is known to be coordinated on the timescales of a few milliseconds (König et al., 1995; Riehle et al., 1997; Butts et al., 2007; Grün, 2009), no observation of sub-millisecond coordination has been reported, nor are there any known mechanisms that enable sub-millisecond synchronization. A high amount of HSEs almost certainly signifies artifacts.

As a first step to mitigate artifacts, after their observation, we tried to identify their physical sources in the experimental recording setup. The first suspect was noise on the reference electrode; the electrode arrays in the macaque Y recording use a common reference wire, which is positioned on the dura (see Materials and Methods). We found many short lived artifacts associated with external events such as switching on a light or closing a door, probably affecting the reference wire. In such interference on the reference wire, the signals on all channels are affected (example for such an event in Common noise artifact example). However, most of the observed HSEs occur only in a small subset of the electrodes. Therefore these events do not originate from the reference electrode. Crosstalk was another potential source of artifacts. It had been known that they tend to happen between electrode shanks at relatively short distances (Nelson et al., 2017). Nevertheless, we observed artifacts also at large distances (Figure 2), suggesting that cross-talk is happening elsewhere. Furthermore, the placement of the individual channels in the connector to the head stage does not seem to explain the spatial patterns of the cross-talk (Figure S2). Therefore, the circuitry inside the headstage is our primary suspect for cross-talk. There the analog signals from many channels are multiplexed, amplified, filtered and converted to digital signals, steps known to be very sensitive to cross-talk (Pérez-Prieto and Delgado-Restituto, 2021; Perez-Prieto et al., 2021). Indeed, in the case of macaque Y, previous recordings had used the analog signal processing headstage ‘Samtec’ and ‘Patient Cable’ (Blackrock Microsystems) (Riehle et al., 2013; Brochier et al., 2018), and after updating it to ‘Cereplex E’ (Blackrock Microsystems) we observed by far fewer artifacts. This is mainly due to the fact that with Cereplex E, digitalization is done at the headstage level, whereas it was done at the amplifier level with the Samtec and patient cables. In the case of macaque L, ‘Cereplex M’ (Blackrock Microsystems) was used. Since it was not possible to further pin down and remove the sources of cross-talk, we decided to develop the post-hoc artifact removal method presented in this paper.

Based on the assumption that artifacts result from linear mixing of signals, we suggested cleaning of data by use of ZCA whitening for artifact removal. This cleaning method works perfectly on simulated data to decorrelate the mixed signals, eliminating the artifacts and recovering the identical signals as in the ground truth data as demonstrated with the cross-talk model. This is because, under the assumption of independent background fluctuations in the ground truth signals, ZCA whitening is exactly the inverse operation of the linear mixture of the ground truth signals with a symmetric mixture matrix, as is done in the model with the cross-talk strength matrix. Application to the Utah array recordings also showed remarkable performance in removing artificial HSEs with high complexity orders, providing a convincing support for the hypothesis that the artifacts in these recordings were generated by electric cross-talk between channels. Besides removing artifacts, ZCA whitening also considerably improved the signal-to-noise ratio of spike waveforms in many channels. This is an expected result of the signal decorrelation by ZCA whitening: the linear mixture of signals by the cross-talk accumulates variances of background fluctuations across channels, and the decorrelation decomposes them into the original variances. Thus, ZCA whitening applied on the Utah array recordings generally improves the signal quality. It does not only remove artifacts, but also reduces background fluctuations in the channels affected by cross-talk, which is beneficial for reliable spike extraction and also successive spike sorting.

Previous studies have taken different approaches for artifact removal. Torre et al. (2016) performed a complete removal of the HSEs with complexities > 1 on the 1/30 ms timescale. This is for sure a brute force approach, but despite that the authors found excess spike synchrony (on the ms scale) for pairs and higher-order synchrony patterns, which revealed a particular spatial organization in the cortex suggested to relate to functional properties. In Brochier et al. (2018) two methods were used to remove artifacts. The first was to define “invalidated waveforms” which were defined by the Plexon Offline Sorter (Plexon Inc, Dallas, Texas, USA) as synchronous potential waveforms involving more than 70% of all channels. This was intended to remove artifacts (e.g., chewing artifacts). These invalidated waveforms were ignored in further analysis including spike sorting. The second step was applied after spike sorting. Here the sorted spike data were controlled on their original time resolution (1/30 ms) for potential occurrences of HSEs using the complexity distribution. HSEs with complexity larger or equal to 2 were removed, as well as spikes occurring within a short time interval around this event (±1/30 ms), which led to a more aggressive removal of HSEs than in Torre et al. (2016). These HSE-removing approaches to artifact removal may damage the data, since the lack of chance synchrony (see Supplementary Figure S1) can be noticed even on longer timescales on the order of several ms. As a result, potentially existing neuronal spike synchrony may be undetected (Oberste-Frielinghaus et al., 2025). Our proposed approach selectively removes the HSEs that stem from problematic correlations between channels, and hence affects the synchrony structure of the original data only minimally. Another previous study (Chen et al., 2022) defined a ‘synchrofact participation’ (SP) ratio as the number of synchronous spike events above chance, derived by a bootstrap method on the complexity distribution. The channels with highest SP were removed iteratively, such that chance levels were re-evaluated after removal of each channel. Thus, instead of individual HSEs, here the whole signal of a channel is removed from the data. Our proposed method provides an alternative approach for artifact removal where no channels need to be removed from the data. For Neuropixels recordings (Jun et al., 2017) artifacts are intended to be removed on the level of the spike sorting. The Kilosort spike sorter (Pachitariu et al., 2024) contains an automatic setting that applies the ZCA whitenting procedure on 32 neighboring channels at a time, successively. However, as we have shown here this should be extended to the application to all channels simultaneously to completely remove the artifacts.

In summary, we showed that artifacts are ubiquitous in multi-electrode recordings. There are multiple possible reasons for their emergence: in the Utah array recordings, they are likely caused by electric cross-talks between the channels, and in the Neuropixels recording likely by the influence of the reference channel in the middle of the electrode. For the detection of the artifacts, we suggest to compare the complexity distribution of the data with their surrogate data at the data sampling resolution. The approach we propose here is to remove artifacts by ZCA whitening. Note that this requires access to the original broadband signal at data sampling resolution. The data cleaning based on ZCA whitening effectively removes artificial HSEs with high complexity orders without removing individual channels, irrespective of the cause of the artifacts. If higher-than-chance complexity events remain after the data cleaning, artifacts originating from common external noise may be present and should also be removed by excluding the affected time periods from further analysis. Artifacts are ubiquitous in multi-electrode neural recordings and their presence can affect results of data analyses. Therefore detection and removal of artifacts is crucial for avoiding false results and ensuring a sound interpretation of the obtained results.

## Supplementary Information

### Complexity distribution after removing all synchronous spikes

We examine the effect of the naive approach to remove all HSEs from the spike trains as proposed by Torre et al. (2016). Therefore we calculate the complexity distribution for the data after all spikes in HSEs are removed and compare this to the surrogates obtained from the data after the removal of the spikes. As to be expected, the distribution for the original data after the removal only contains entries for complexities of zero and one. However in the surrogates we find complexity up to six in the surrogate data, concluding that the data after removal significantly lacks synchrony since a certain amount of HSEs is expected. This shows that the naive approach should not be taken to remove artifacts.

### Connector mapping versus array mapping

In search for causes of strong band-passed signal correlations between channels, we examine the spatial distribution of the channels showing strong correlations on two spatial maps of channels: one on the level of the electrode arrays and the other the head stage connectors. In the case of macaque Y, the relative positions of channels are almost identical between the two maps, and hence we cannot conclude on which level the cross-talk is localized. In the case of macaque L, however, the spatial distribution of highly correlated channels is largely different between the array mapping and the connector mapping, indicating that the cause of the high correlations is localized on the level of the electrode arrays.

### Comparison between empirical and surrogate HSE index

To examine how large the HSE indices obtained from real data are in comparison to those from independent spike trains with matched firing rates, we plot the empirical pairwise HSE indices in the macaque Y data against the respective surrogate HSE indices obtained from time shifted surrogate data (Figure S3a; the mean over 200 surrogates is taken for each channel pair). The majority of channel pairs show larger empirical HSE index values than the respective surrogate values. This could reflect a bias originating from the definition of the HSE index that artifacts in channels with small numbers of spikes can be over-represented in the index. Hence, we next focus only on channels with firing rates greater than 1 spikes/s (Figure S3b). While a certain amount of channel pairs with large empirical HSE indices were screened out by this conditioning, we still see a considerable number of channels with extremely large empirical HSE indices compared to the surrogates, which are most likely the ones suffering from the cross-talk. After applying our proposed channel exclusion methods (Figure S3c and d), those channel pairs are screened out and the remaining ones show empirical HSE indices comparable to the surrogate indices, meaning that the remaining channels have as many HSEs as in independent spike trains. The mean of the empirical HSE indices over channel pairs is still slightly higher than the mean of the surrogate HSE indices, which is likely due to other causes of artifact than cross-talk. One of such causes is common external noise across channels. Spike-like events generated by such noise typically show waveforms dissimilar to real spike waveforms, and hence can be effectively excluded by spike sorting. To check whether that is actually the case, we plot the HSE indices for SUA pairs in the same manner as before (Figure S3e-g). Again, after applying our proposed methods, SUA pairs show only as many HSEs as in the surrogates, and in this case of SUA pairs, the mean empirical HSE index is much more consistent with the mean surrogate HSE index than in the case of channel pairs.

### Common noise artifact example

The data of macaque L contained highly synchronized activity in the data lasting for about 500 ms (see Figure S4, bottom), which is likely due to strong common noise contaminating all the channels equally. Such common noise can enter the system via the reference electrode. We identified this burst of synchronous spike activity as artifacts since these synchronous spikes also still exist on the sampling rate resolution (top). It introduced in the complexity distribution a huge amount of events with complexities up to 35. Therefore, we removed this piece of data (1000 ms) before any further analysis. We did not detect such events in the data from macaque Y.

---

**Figure S1.**
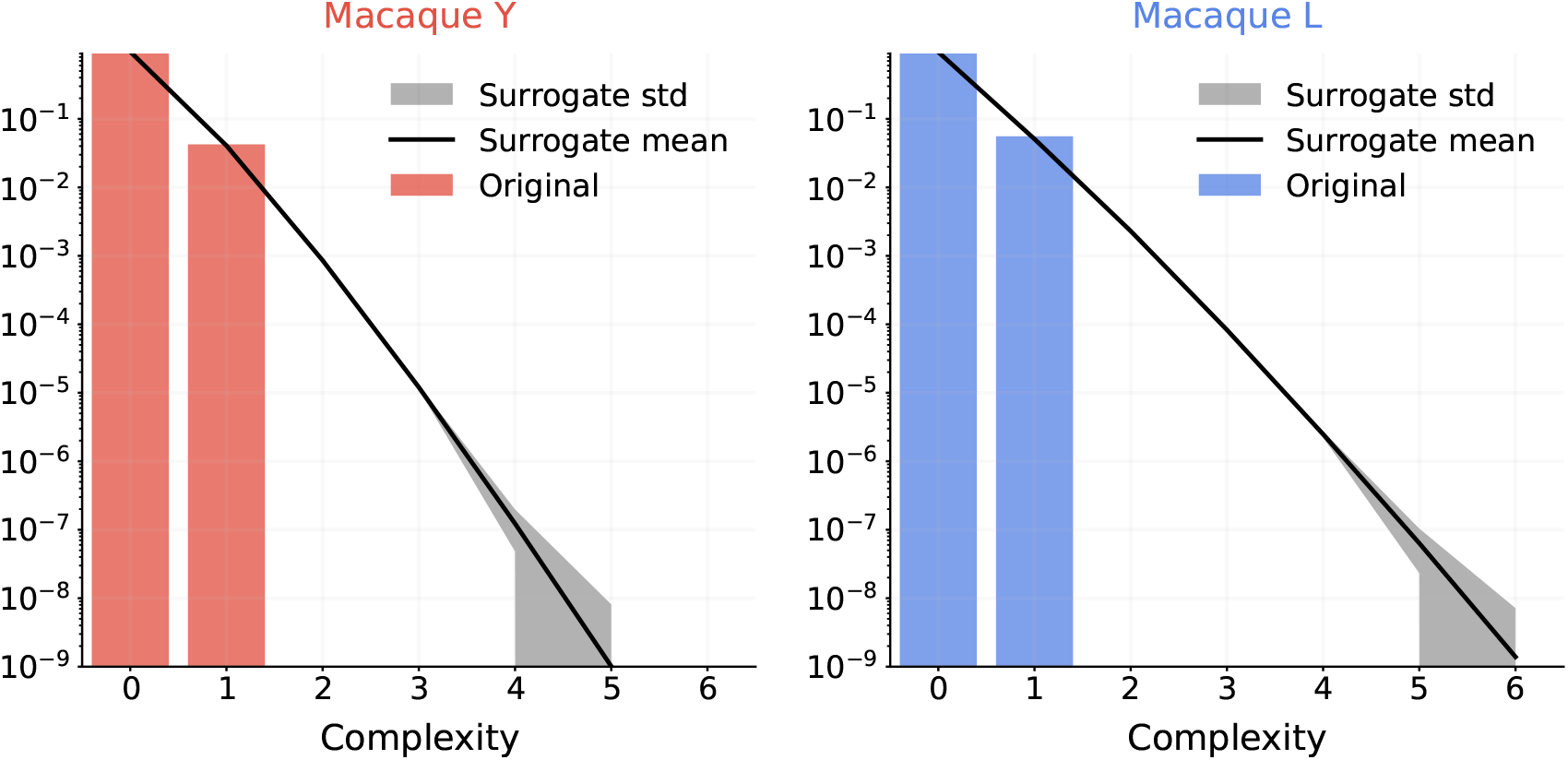
Complexity distribution after removing all synchronous spikes, Complexity distribution of the original data (colored bars), and the mean and standard deviation (line and shade, respectively) for the complexity distribution of the respective surrogates, 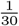ms bin size.

**Figure S2.**
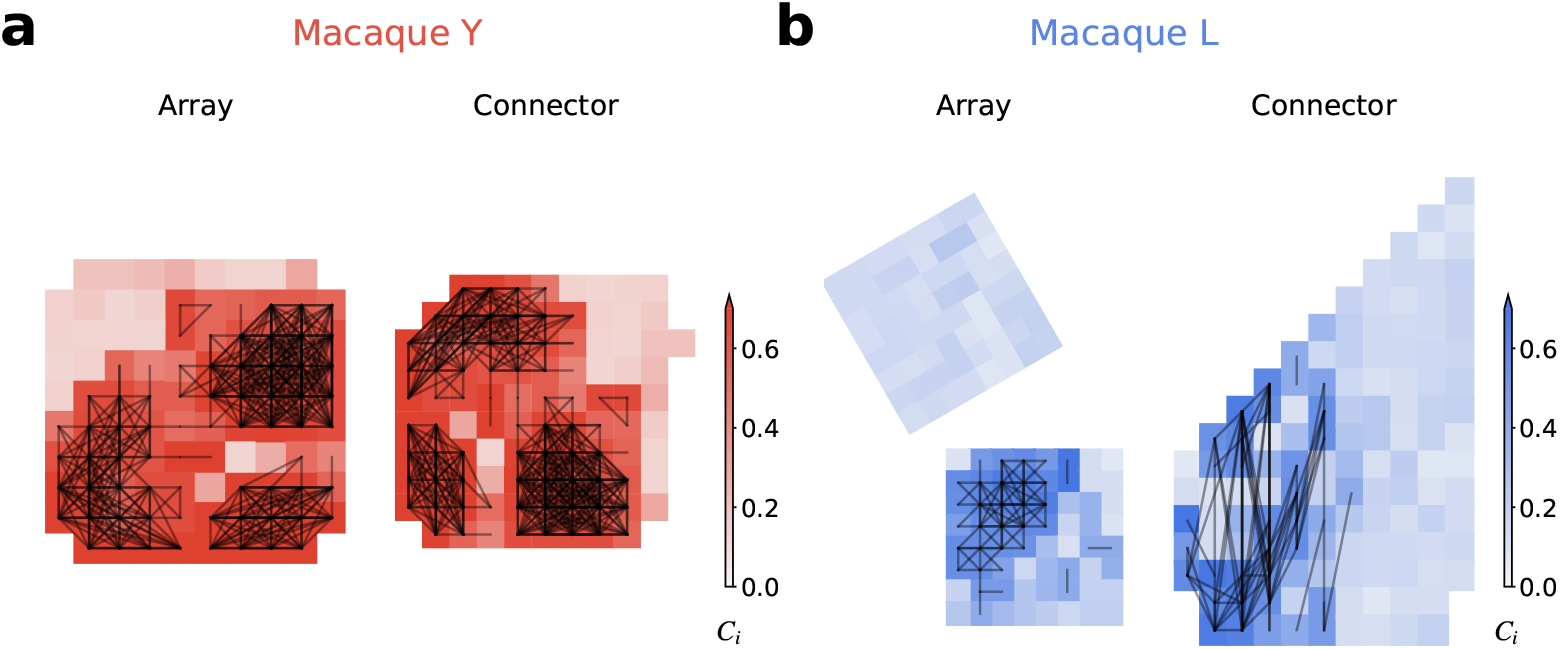
Comparison of spatial distribution of maximal band-passed signal correlation per channel (*C*_*i*_) on the electrode array layout (left) and the headstage connector layout (right) for (a) macaque Y and (b) macaque L. Black lines connect electrode pairs with pairwise band-passed signal correlations *c*_*i,j*_ > 0.4.

**Figure S3.**
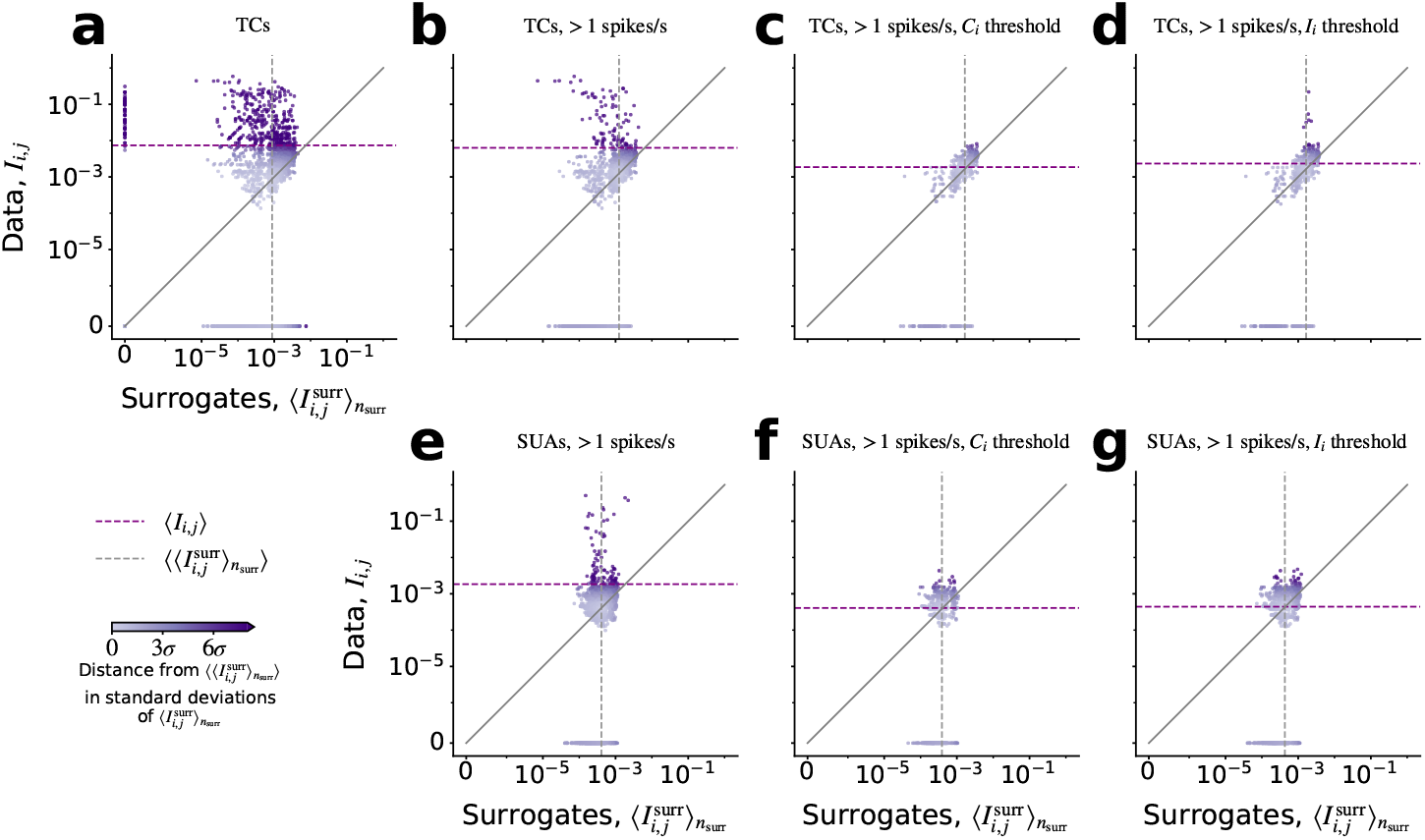
Comparison of the pairwise HSE index *I*_*i,j*_ in the experimental data and the average of the corresponding surrogate data 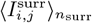, each point corresponding to one channel pair. Mean of each variable is shown with a dashed line. The diagonal line indicates the identity between the two measures. The color shading represents the difference from the theoretically random data, at the point where the mean surrogate index 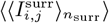 crosses the diagonal; large deviations from this point are indicative of above-chance HSEs. The different panels show the results for the same session based on (a-d) all threshold crossings (TCs), or (e-g) spike sorted single units (SUAs). The different columns show different removal methods: (b, e) threshold on the firing rate, (c, f) threshold on firing rate and maximum cross-correlation *C*_*i*_, and (d, g) threshold on firing rate and global HSE index *I*_*i*_.

**Figure S4.**
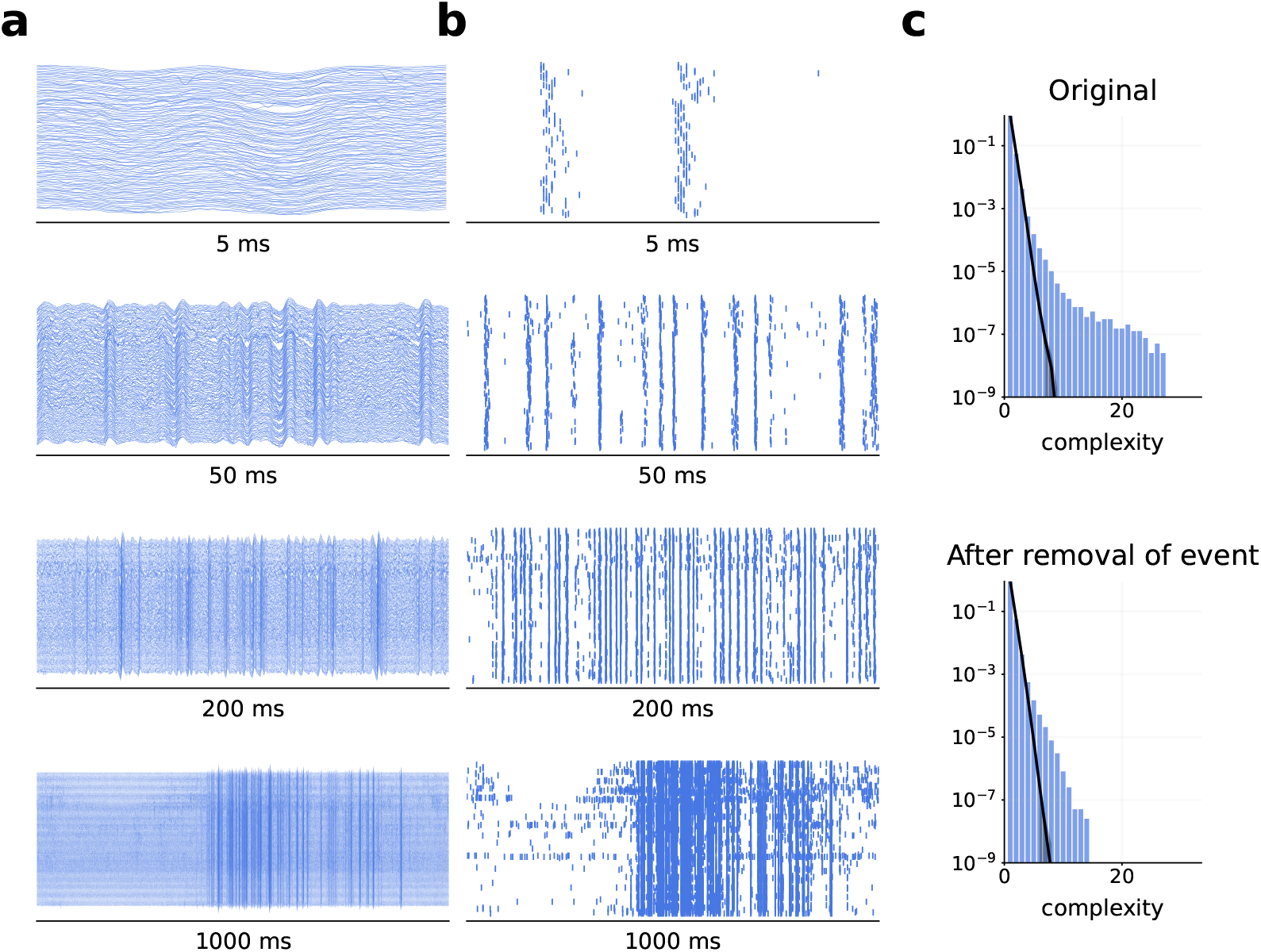
Burst artifact in macaque L. The left column, bottom shows the whole burst artifact, that expresses a lot of artifacts (on sampling rate of 30 kHz) one after the other over a period of about 500 ms. The plots above show higher resolutions of that event, and on the top the total display shows piece of the data lasting 5 ms. On the right top, the complexity distribution of the whole data set is shown, below after the complete removal of this burst event.

## Footnotes

1 Note that, for simplicity, this formalism does not consider “noise” spikes which arise from the random fluctuations of the background signal that happen to cross the threshold by chance.

## Notes

### Competing Interest Statement

The authors have declared no competing interest.

### Summary of Updates

Revision of discussion Changed order of sections Additional information about the setup

